# The potato cyst nematode *Globodera pallida* overcomes major potato resistance through selection on standing variation at a single locus

**DOI:** 10.1101/2025.07.23.664352

**Authors:** Arno S. Schaveling, Dennie M. te Molder, Paul Heeres, Joris J.M. van Steenbrugge, Stefan J.S. van de Ruitenbeek, Casper C. van Schaik, Sven van den Elsen, Geert Smant, Mark G. Sterken

**Affiliations:** Laboratory of Nematology, Wageningen University & Research, Droevendaalsesteeg 1, 6708 PB Wageningen, the Netherlands; Averis Seeds B.V., Valtherblokken-Zuid 40, 7876 TC Valthermond, the Netherlands; Department of Genetics, University Medical Center Utrecht, Utrecht University, The Netherlands

**Author notes:** Corresponding author Mark Sterken Laboratory of Nematology Wageningen University and Research Droevendaalsesteeg 1 6708PB, Wageningen.

**Keywords:** *Globodera pallida*, virulence, standing variation, resistance, potato, effector, *GpaV_vrn_*

## Abstract

*Globodera pallida* poses a major threat to potato production, with management strategies primarily relying on genetic resistance. However, reports from multiple locations in Western Europe indicate a steady increase in virulence levels among field populations, raising major concerns about *G. pallida* control. The evolutionary mechanisms driving this rise in virulence are poorly understood.

To investigate this, we analysed the propagation of thirteen recently isolated field populations on thirty commercial potato varieties over four independent PCN resistance tests. Our findings indicate that (1) the genetic basis of resistance in potatoes is small, with the major resistance conferred by *GpaV* from *Solanum vernei*, and (2) the wide application of *GpaV_vrn_* has led to continuous selection on standing genetic variation in *G. pallida* field populations. To map virulence, we propagated two field populations on a *GpaV_vrn_*-resistant variety for four consecutive generations. High-coverage whole-genome sequencing of each generation revealed that *GpaV_vrn_*-mediated selection acted on a single locus of a newly assembled *G. pallida* Rookmaker reference genome. Examination of this virulence-associated locus identified *Gp-pat-1* as a candidate gene. Silencing *Gp-pat-1* increased *G. pallida* virulence on a *GpaV_vrn_*-resistant potato variety but had no effect on nematode virulence on a susceptible variety. Thereby classifying *Gp-pat-1* as an avirulence gene and confirming its role in the breakdown of *GpaV_vrn_*-resistance.

These findings demonstrate that negative selection on the *Gp-pat-1* avirulence allele by *GpaV_vrn_*-mediated resistance is driving the emergence of virulence in the *G. pallida* field populations used in our selection experiments. Our results therefore strongly suggest that selection on *Gp-pat-1* is likely underlying the current outbreak of *GpaV_vrn_*-resistance breaking populations. Our results provide a foundation for the development of molecular diagnostic tools to monitor virulence in field populations, to enhance understanding of resistance breakdown, and to inform the sustainable deployment of resistances in the field.

**SIGNIFICANCE STATEMENT:** In Western Europe, management of the potato cyst nematode *Globodera pallida* primarily relies on genetic resistances in potato plants. However, resistance-breaking populations are emerging across Western Europe. Here, we identify a single resistance locus shared by all tested resistant potato varieties. We also identify a corresponding virulence locus on a newly assembled *G. pallida* reference genome and identify a gene within this locus that contributes to virulence. Our findings provide critical insights into the selective pressure acting on this plant-parasitic nematode, the emergence of virulence over time, and the molecular mechanism underlying resistance breakdown. These results provide much needed insights into a highly adapted soil-borne plant-pathogen.

## INTRODUCTION

The potato cyst nematode (PCN) species *Globodera pallida* and *G. rostochiensis* threaten the European potato production (Hockland et al., 2012; Jones et al., 2013). PCN infestations cause symptoms that include stunted growth, leaf yellowing, and reduced tuber size, leading to substantial yield losses (Price et al., 2021). The use of resistant potato cultivars has long been the cornerstone of PCN management (Gartner et al., 2021). Following a major *G. pallida* outbreak in the 1990s, resistant varieties that carried resistances from wild *Solanum* species were introduced to control *G. pallida* (Grenier et al., 2020). However, the deployment of these resistances has exerted strong selection pressure on *G. pallida* populations. Over the last decade, resistance-breaking *G. pallida* populations have become a matter of great concern (den Nijs & van Heese, 2019; Mwangi et al., 2019; Niere et al., 2014). Despite previous research (Eoche-Bosy, Gauthier, et al., 2017; Eoche-Bosy, Gautier, et al., 2017; Lechevalier et al., 2025; Varypatakis et al., 2020), the genetic basis of potato resistance and the mechanism through which *G. pallida* overcomes this resistance remain elusive.

Potato resistances typically affect PCN feeding site formation. As PCN are obligate sedentary endoparasites, their survival relies on the development of a syncytium. The syncytium is formed through the dissolution of plant cell walls and subsequent fusion of neighbouring protoplasts (Goverse et al., 2000). Mature syncytia can consist of hundreds of fused host cells, supplying nutrients critical for nematode development and maturation. Disrupting syncytium formation or functioning are effective mechanisms of resistance (Goverse & Smant, 2014). Resistant potato varieties generally induce strong necrosis around the head of sedentary juveniles to prevent syncytium initiation, or around the syncytium to hinder syncytium development, arresting juvenile development (Fournet et al., 2018; Rice et al., 1987; Varypatakis et al., 2020). Although numerous sources of potato resistance against *G. pallida* have been mapped [(Gartner et al., 2024; Leuenberger et al., 2025); and reviewed in Gartner et al. (2021)], the genetic basis of potato resistance in commercial cultivars has yet to be defined.

Resistance-breaking *G. pallida* field population have been reported in Emsland, Germany (Niere et al., 2014) and northeastern Netherlands (den Nijs & van Heese, 2019; Grenier et al., 2020). These populations have been reported to break through resistances in cultivars that were previously identified as resistant based on standard *G. pallida* testing populations. Previously, many studies have explored phenotypic and genetic adaptations of *G. pallida* to resistant potato cultivars in experimental settings (Eoche-Bosy, Gauthier, et al., 2017; Eoche-Bosy, Gautier, et al., 2017; Fournet et al., 2018; Fournet et al., 2013; Lechevalier et al., 2025; Mwangi et al., 2019; Varypatakis et al., 2020). However, the genetic basis and molecular mechanisms underlying virulence on resistant potato varieties are still poorly understood.

In this study, we aimed to identify the genetic basis of *G. pallida* resistance in commercial potato varieties and resolve the cause of increasing virulence in Dutch *G. pallida* field populations. We discovered the source of resistance, *GpaV_vrn_*, is shared by all resistant potato cultivars tested. We tested the hypothesis that *G. pallida* virulence is caused by selection on standing variation already present at the time of introduction of the resistance varieties. We associated virulence with a single locus on the newly assembled *G. pallida* reference genome. We applied three criteria to identify putative resistance-breaking effector genes within the avirulence locus and pinpointed *Gp-pat-1* as a candidate gene for avirulence on *GpaV_vrn_*. Silencing *Gp-pat-1* in juveniles significantly increased virulence on resistant potatoes, confirming its role in the breakdown of *GpaV_vrn_*.

## Materials and methods

### G. pallida populations

Thirteen populations of *G. pallida* were collected from infection foci in fields with PCN resistant potato varieties. These populations were gathered from 2011 – 2015 and originate from the North-East of the Netherlands. All populations were propagated on the susceptible potato variety Desirée and confirmed to consist exclusively of *G. pallida* through species-specific PCR testing. The Dutch population Pa_3_-E400 (Rookmaker) and the European population Pa_3_-Chavornay, which are commonly used for selecting PCN resistant breeding material, were included for reference. The Pa_2_-D383 and the Rookmaker populations were used for genomic comparisons.

### Potato varieties

Thirty one different resistant and widely used commercial potato varieties were used across four standard PCN resistance tests for quantifying reproduction of potato cyst nematodes on potato (EPPO, 2013). The presence of the *GpaV_vrn_* resistance within these potato varieties (except the susceptible reference cultivar Desiree) was confirmed with a genetic marker by breeding companies.

### General notes on data analysis

All tests and analyses were conducted in “R” (version 4.1.0 win x64) using Rstudio (version 1.4.1717; R Core Team, 2013; Team, 2015). Only for the alignment of the sequencing data and variant calling other software was used. In R, the *tidyverse* packages, especially *ggplot2* and *dplyr* were used for data processing in general and generation of most of the figures. Other, specific packages used are mentioned at the relevant sections.

All scripts and underlying phenotypic datasets are available through gitlab (https://git.wur.nl/published_papers/Schaveling_2025_Pallifit_virulence). The data of standardized PCN resistance test and the small container tests are included in the supplementary data. The Rookmaker genome, together with the underlying DNA and RNA sequencing data was deposited at the European Nucleotide Archive (ENA; PRJEB91928). The DNA sequencing data of the selection experiment was deposited at the ENA (E-MTAB-15408)

### Standard PCN resistance tests

Four standard PCN resistance tests were conducted according to the EPPO standard protocol (EPPO, 2013), the resistance tests are ordered based on appearance in results section.

The first standard PCN resistance tests were executed in 2019 at the NAK (Emmeloord, the Netherlands). The goal of this PCN resistance test was to compare the multiplication of two previously tested virulent *G. pallida* populations (i.e., AMPOP02 and AMPOP13), and three newly collected field populations (i.e., 2017.Pa.2018, 2017.Te.2018, and 2017.dC.2018), with two standard *G. pallida* populations (Pa_3_-E400 and Pa_3_-Chavornay) on sixteen commercial potato varieties (i.e., Alcander, Allison, Altus, Ardeche, Arsenal, Avarna, Avito, Axion, Basin Russet, Desiree, Festien, Innovator, Libero, Lugano, Seresta, and Supporter). The PCN resistance test included four replicates for each combination of potato variety and nematode population.

The second standard PCN resistance test was executed in 2017 at the HLB (Wijster, the Netherlands). Herein, the multiplication of nine field populations of *G. pallida* (i.e., AMPOP01, AMPOP02, AMPOP03, AMPOP06, AMPOP09, AMPOP10, AMPOP13, AMPOP15, and AMPOP18) was tested on fifteen commercial potato varieties with strong *G.* pallida resistance according to the national cultivar list of the Netherlands (i.e., Altus, Ardeche, Arsenal, Avarna, Avito, Axion, Basin Russet, Festien, HZD 06-1249, Innovator, Libero, Seresta, Supporter, and VD 07-0289) and the reference cultivar Desiree. Each combination of potato variety and *G. pallida* population was replicated three times. To this end, single potato eye plugs were planted into 2 litre pots containing approximately 2000 grams of soil inoculated with 5 juveniles per gram of soil. The pots with the potato plants were maintained in a greenhouse under natural light conditions from February to May. One hundred days after the start of the experiment the soil was allowed to dry slowly. The cysts were extracted from the dry soil by elutriation, whereafter the number of cysts per pot and the number of living juveniles and eggs per cyst were quantified. The number of living larvae and eggs were counted in triplicate.

The year after, in 2018, a third PCN resistance test was conducted at the HLB with seven *G. pallida* populations (i.e., AMPOP01, AMPOP02, AMPOP03, AMPOP06, AMPOP09, AMPOP10, and AMPOP18) and sixteen commercial potato varieties (i.e., Actaro, Avatar, Aveka, Aventra, BMC, Desiree, Festien, Merenco, Novano, Saprodi, Sarion, Sereno, Seresta, Simphony, Stratos, and Vermont) using the same protocol.

The fourth PCN resistance test was conducted at the NAK in 2019. The goal of the fourth PCN resistance test was to quantify an increase in multiplication rate of five consecutive generations of two *G. pallida* field populations selected on Seresta (i.e., AMPOP02 and AMPOP10) on a set of six potato varieties (i.e., Avarna, Desiree, Festien, Innovator, Libero, and Seresta). For this experiment, each combination of potato variety and *G. pallida* population was replicated three times. Unfortunately, there were four errors in the quantification of the AMPOP10, especially in Seresta generation 1 (AMPOP10DS_2016). The dilutions were not administered correctly, leading to extreme amounts of nematodes per cyst (>>500) in the Desiree, Seresta, Festien, and Avarna counting. The cyst counting data was not affected, therefore we could reconstruct the dilution mistakes (one two-fold and three four-fold mistakes) by relying on the propagation in AMPOP10D and AMPOP10DS2 (as the within-population number of cysts per larvae per variety were stable).

### Data normalization of standardized PCN resistance tests

To make integrated analysis possible, the second and third standard PCN resistance tests were normalized based on the three potato varieties that were measured on seven identical *G. pallida* field isolates in each year. For each year, the average Pf/Pi, number of cysts, and number of larvae was calculated over these potato varieties and *G. pallida* populations and the ratio over the years was calculated to provide a normalization constant, by

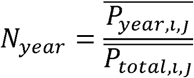

where N_year_ is the normalization constant, and 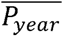 is the average phenotype over potato varieties *i* (Desiree, Festien, and Seresta) and *G. pallida* populations *j* (AMPOP01, AMPOP02, AMPOP03, AMPOP06, AMPOP09, AMPOP10, and AMPOP18). 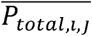 is the total average over the two years. The phenotypes were normalized by

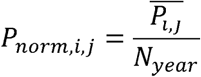

Where 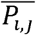 is the trait average over three replicates of potato variety *i* (one of the remaining 28 potato varieties), and *G. pallida* population *j* (one of 10 populations).

### Data analysis of standardized PCN resistance test

Differences between groups were determined using two-sided t-tests as implemented in the R-package ggpubr (Kassambara, 2018).

A cluster analysis was conducted to compare the potato varieties and *G. pallida* populations. A Euclidian distance matrix was calculated based on the averaged normalized Pf/Pi values per potato variety per *G. pallida* population; using the *dist* function in “R”. These were clustered using the *hclust* function in “R”. To further test the clusters, a pairwise Tukey-test was conducted, for which the p-values were corrected for multiple testing using Benjamini-Hochberg correction as implemented in *p.adjust* function in “R”. For plotting, the p-values were log_10_-transformed, the minimum untransformed p-value was capped at p = 1*10^-10^.

Analyses of variance were done using a PERMANOVA, for which we used the *adonis2* function of the *vegan* package (Oksanen et al., 2013). The PERMANOVA model for analysis of the contributions factors to virulence was ran on the normalized propagation data (Pf/Pi) using

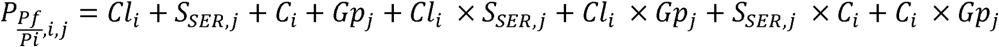

where *P_Pf/Pi_* is the normalized propagation of *G. pallida* population *j* (one of nine virulent populations) on potato variety *i* (on of 28 potato varieties). *Cl* is the cluster the potato variety belongs to (Cl_SER_, CL_FES_), S_SER_ is the mean Pf/Pi on Seresta-cluster potato varieties. *C* is the potato variety (one of 28), *Gp* is the *G. pallida* population (one of nine). The significances were calculated using 10,000 permutations. It should be noted that this model excluded the two varieties that were not in the Seresta and Festien cluster (Desiree and Aventra). Exclusion reduced the amount of variance captured by cluster (as they show very high replication of the populations).

### Potato pedigree analysis and SNP data

Potato pedigrees were obtained via the potato pedigree database (Van Berloo et al., 2007). The pedigree contains information on 28/31 potato varieties tested in the presented experiments (HZD 06-1249, Novano, and VD 07-0289 were not included in the database). The whole database was downloaded and initially processed using custom scripts to map the resistance progenitors of the potato varieties. Subsequently, by hand, the progenitors were annotated for underlying wild potato sources, focusing on sources with known resistance contributions: *Solanum multidissectum*, *S. oplocense*, *S. sparsipilum*, *S. spegazzinii*, *S. tarijense, S. tuberosum* ssp. *andigena*, *S. vernei* (Gartner et al., 2021; Rouppe Van der Voort et al., 2000; van der Voort et al., 1998). Subsequently, these sources were mapped back to current varieties.

For 22 of the 31 potato varieties tested in the presented experiments additional SNP data was generated by the breeding companies Averis, Agrico and HZPC. The SNP data was based on the SolSTW SNP array platform (Vos et al., 2015), and the potato PGSC v4.03 pseudomolecules (Sharma et al., 2013; Xu et al., 2011). The data were analysed by graphical mapping, similar to (van Eck et al., 2017). We used the susceptible variety Desiree and the resistant variety Innovator for *GpaV_vrn_* and on the susceptible variety Desiree and the resistant variety Seresta for *Gpa6*.

### Small container PCN resistance tests

As a service for potato producers, small container tests for PCN resistance in potato are commercially offered at the HLB to predict the performance of potato varieties on infection foci of potato cyst nematodes. These small container tests for PCN resistance in potato are conducted in 55 ml transparent plastic containers harbouring a small tuber and soil inoculated with 300 juveniles of *G. pallida* per container. A test includes eight replicates for each combination of potato variety and *G. pallida* population. The containers are kept at 20°C in the dark for 8-12 weeks, after which the number of cysts visible on the root system through the walls of the container are counted. A database of anonymized results of all small container PCN resistance tests commercially executed from 2016 – 2021 by HLB was made available to this study. Notably, multiplication rates of nine of the *G. pallida* populations mentioned above (i.e., AMPOP01, AMPOP02, AMPOP03, AMPOP06, AMPOP09, AMPOP10, AMPOP13, AMPOP15, and AMPOP18) on thirteen potato varieties (i.e., Avito, Supporter, Festien, Altus, Merenco, Saprodi, BMC, Avarna, Sarion, Seresta, Axion, Novano, Desiree) in small container tests were also in the database. More importantly, similar data of 125 infestation foci of *G. pallida*, which were recently sampled on farms and tested on the potato varieties Desiree, Seresta, and Festien in small container tests, were retrieved from the database for further analyses.

### Data analysis of the small container tests

For the correlation analysis of the small container tests versus the standardized container tests, the mean relative susceptibility values (determined versus Desiree) of both tests were used. The analysis was conducted per *G. pallida* population. We had data for comparing the nine virulent populations. A Pearson correlation was calculated as well as the significance of the correlation. To determine how well virulence levels on resistant potato varieties were estimated as compared to standardized container tests, we also analysed the correlation without including Desiree. The regression line of that correlation showed how well virulence levels were estimated.

The between variety differences (Desiree, Seresta, and Festien) were calculated using a t-test on the relative susceptibility values.

### Selection for virulence by GpaV_vrn_

Two *G. pallida* populations with suspected virulence, designated AMPOP02 and AMPOP10, were used to further select for virulence on the *GpaV_vrn_* resistant potato variety Seresta in pot experiments. To this end, a maximum of 10,000 eggs was inoculated into two litre pots containing a 2.2 kg mixture of silver sand, kaolin, hydro grains and nutrients. To each of the pots, a potato piece containing a single shoot was added. To obtain sufficient starting material, prior to the selection experiment, AMPOP02 was propagated on the susceptible variety Desiree for two consecutive generations, resulting in AMpop02D_2_ (**Figure 3A**). AMPOP10 was propagated on Desiree for one generation, resulting in AMpop02D_1_.

The selection experiment started in 2015, going through one generation per year, leading to fourth-generation selection populations at the end of 2018. For each generation we aimed to save cysts to later use for DNA extraction. Since all of the yield from the first round of selection on Seresta (S_1_) was needed for inoculation of the second generation, together with inoculating the second generation, we again inoculated the unselected starting populations on Seresta, to reproduce the S_1_ (**Figure 3A**). It should be noted that the S_1_ populations were independently derived starting from the same Desiree propagation.

### DNA isolation and sequencing of selected G. pallida populations

DNA of the *G. pallida* selection experiment was isolated using previously published protocols (van Steenbrugge et al., 2023). In short, J2 larvae collected after counting were frozen and stored at -80°C. These were lysed (Holterman et al., 2006) and DNA was isolated using phenol/chloroform/isoamyl alcohol. DNA quantities were determined using Qubit Fluorometer (Invitrogen). Because we were dependent on how well the material was preserved after counting not all samples provided DNA of sufficient quantity and/or quality, 45 out of 60 samples were successfully sequenced.

The DNA was sequenced at BGI Genomics (China) using BGISEQ-500 with a median output of 295.4*10^6^ reads of 100 bases (paired) per sample. Upon arrival the samples, integrity and purity were tested by gel-electrophoresis. Per sample 1 µg of DNA was fragmented by Covaris and fragments in the size ranges of 150-250 bp were selected which were subsequently quantified by Qubit. Fragments were end-repaired and 3’ adenylated prior to adaptor-ligation to the 3’ end. The fragments were amplified by PCR and purified with the Agencourt AMPure XP-Medium kit. DNA was quantified by Agilent 4200 TapeStation. Double stranded PCR products were circularized, and the circular DNA was used as input for sequencing in the BGISEQ-500 platform. FASTQ files were deposited at the ENA (E-MTAB-15408)

### Data analysis of the GpaV_vrn_ selection experiment

The data of the fourth standard PCN resistance test was analysed for three phenotypes that could be relevant for virulence: the propagation (Pf/Pi), the relative susceptibility versus Desiree (RS in percentage), and the number of eggs/juveniles per cyst. Due to the amount of propagation achieved, we had material to test consecutive selection generation 2, 3, and 4 for both AMPOP02 and AMPOP10. We did test a non-selected and a first-generation Seresta selected population, as these were not directly related to generation 2, 3, and 4.

Within variety comparisons were conducted using Kruskal Wallis tests. To test for the contribution of generations of selection a PERMANOVA was conducted on generation 1, 2, 3, and 4 for AMPOP02 and AMPOP10 separately using the model

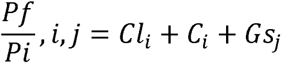

where *Pf/Pi* is the propagation of *G. pallida* population *j* (either AMPOP02 or AMPOP10) on potato variety *i* (one of 6 potato varieties), *Cl* is the cluster the potato variety belongs to, *C* is the potato variety, *Gs* is the generation of Seresta selection of the *G. pallida* population (1 – 4). The significances were calculated using 10,000 permutations. Note that neither generation 1 population was directly related to generation 2 – 4. However, this generation was selected based on the same unselected material used to establish generation 2 – 4, therefore we think it was prudent to include this generation in the analysis.

### Broad-sense heritability analysis

Broad sense heritability was calculated for the potato varieties Desiree, Seresta, Festien, Avarna, Libero, and Innovator for the fourth standard PCN resistance test on field populations, the selection experiment, and the small container tests (only Desiree, Seresta, and Festien). The heritability was determined independently for the AMPOP02 and the AMPOP10 selected populations. We used the equation

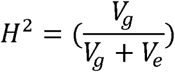

where *H^2^* is the (broad sense) heritability, *V_g_* is the variance explained by population, and *V_e_* is the residual variance. The *V_g_*and *V_e_* components were determined using a REML model as implemented by lme4: RS ∼ 1 + (1|population) (Bates et al., 2015). It should be noted that depending on the populations used the interpretation of *H^2^* varies. Namely, in the standardized PCN resistance test on various populations as well as the small container tests, it is a measure of variance captured by populations for which we did not include their genetic relations. It was therefore probably a less accurate estimation of the true heritability. The *H^2^*for the selection experiments was probably the most reliable estimator of the heritability, as the populations were directly related.

### Promethion sequencing and assembly of G. pallida Rookmaker genome

DNA isolation of the *G. pallida* Rookmaker population was performed as described before (van Steenbrugge et al., 2023). Long-read DNA sequencing was done by USEQ (The Netherlands). Raw Oxford Nanopore promethION reads (ERR15205749) were corrected to merge haplotypes using the correction mode in Canu (Koren et al., 2017), by reducing the error rate to a maximum of 14.4% and the corrected coverage to a minimum of 400.

Genome assembly and polishing was conducted as described before (van Steenbrugge et al., 2023). In brief, using wtdgb2 v2.5 (Ruan & Li, 2020), multiple initial genome assemblies were generated based on the corrected Nanopore reads while manually refining the parameters minimal read length, k-mer size, and minimal read depth. These parameters were optimised to generate an assembly close to the expected genome size of *G. pallida* (110 Mb). After optimisation, for *G. pallida*, a minimum read length cut-off of 4,000, minimal read depth of 12, and a HPC k-mer size of 19 was used. Remaining haplotigs were pruned from the assemblies using Purge Haplotigs v1.1.2 (Roach et al., 2018). Based on the histogram, low read depth cutoff parameter was set at 75, the low point between haploid and diploid peak was determined at 375, and the read depth high cutoff was set at 650. Contigs were improved using FinisherSC v2.1 (Lam et al., 2015) at default settings and scaffolded using LongStitch v1.0.4. with a HPC k-mer size of 32 and a window size of 500 (Coombe et al., 2021).

The scaffolded genome was filled with RagTag v2.1.0 (Alonge et al., 2022) using the closely related *G. pallida* D383 genome (van Steenbrugge et al., 2023) the resulting assembly was polished with Nanopore reads by Medaka v1.4.4 (https://github.com/nanoporetech/medaka), model r941_min_high_g360, followed by five iterations of polishing with Pilon v1.24 (Walker et al., 2014) using Illumina HiSeq reads from the Rookmaker population. Gene annotations were predicted using Braker v2.1.6 (Brůna et al., 2021) with a modified intron support of 0.51, aided by RNAseq data of parasitic and pre-parasitic Rookmaker life stages. The data used includes: Rookmaker samples of E-MTAB-11646 (Zheng et al., 2022), life-stage samples of PRJEB2896 (Cotton et al., 2014) and our own generated RNAseq of mixed stages of Rookmaker (ERR15277786). We also used the protein data base of OrthoDB (v10; Kriventseva et al., 2018). The genome is deposited at the European Nucleotide Archive (ENA) under PRJEB91928.

### Sequence alignment and variant calling of the GpaV_vrn_ selection experiment

The reads were aligned to the newly assembled Rookmaker genome and the previously constructed *G. pallida* D383 genome (van Steenbrugge et al., 2023) using bwa-mem2 (v2.2.1; Vasimuddin et al., 2019) with the default parameters. Duplicate reads were marked using Samtools MarkDups (v1.14; Danecek et al., 2021). Variants were called using Bcftools (mpileup & call, v1.14; Danecek et al., 2021) with the default parameters except for a maximum number of 1500 reads per position (-d), an INDEL threshold of 1500 reads (-L), and a minimum mapping quality of 20 (-q) after mismatch correction (-C50). The initial round of vcf filtering was performed using Bcftools’s vcfultils.pl with the default parameters except for a minimum read depth (-d) of 10 and minimum alt allele depth (-a) of 5. For the remaining calls the Alternative Allele Fraction (AAF) was computed by dividing the Allelic Depth (AD) of the most common alternative allele (across all samples) by the combined AD of the reference allele and most common alternative allele (bi-allelic depth). Single calls were marked as missing if the fraction of bi-allelic reads was less than 0.95 or if the bi-allelic depth was less than 5. The resulting variant matrix spanned a continuous range from 0 to 1, which was better able to represent the allele fractions present in heterogeneous populations. The pipeline was orchestrated by Snakemake (v6.8.0; Mölder et al., 2021) running on python (v3.7.8; https://git.wur.nl/stefan.vanderuitenbeek/dnaseq_variant_calling_snakemake_pipeline).

A second round of filtering was applied to the variant matrix based in part on statistics generated by vcftools (0.1.16; Danecek et al., 2011) including: a minimum quality of 30, a minimum and maximum mean read depth of 30 and 300 respectively, at least 10 alternative reads, single nucleotide polymorphism variants only (as these are more reliably called), mean alternative allele frequencies between 0.05 and 0.95, and no missing calls. These filtering steps led to the detection of 1,216,526 variants on the Rookmaker genome and 1,132,495 variants on the D383 genome.

### Analysis of genetic variation of the GpaV_vrn_ selection experiment

To understand the genetic relation between the samples, the Rookmaker variants were analysed in a principal component analysis using *prcomp* with the parameter scale. = TRUE. The first four axes (capturing 96.9% of variance) were graphically analysed.

The generational model applied on both sets of variants was a linear model, where the assumption was that in the range where we were measuring the allele frequency increased linearly per generation. This assumption would not be true at either very low or very high relative susceptibility. Since we measured relative susceptibilities within the range of 16.7 till 80.3 on Seresta, it is likely that our measurements are in the range where allele frequencies increase in a more-or-less linear relation with generation (Schouten, 1993). Therefore, we used

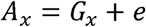

where *A* was the allele frequency in sample *x* (one of 23 from AMPOP02 and one of 22 from AMPOP10), *G* was generation of sample *x* (0, 1, 2, 3, 4, or 5) and *e* is the error-term. The obtained significances were Bonferroni corrected. This model was run separately for AMPOP02 and AMPOP10.

The locus containing the virulence gene was determined based on changepoint analysis, using the changepoint package in R (Killick & Eckley, 2014). The association outcomes of AMPOP02 and AMPOP10 were combined. We used the associated variant density to identify the most SNP-dense associated region on scaffold 28.

The Venn-diagram illustrating the overlap in significant variants derived from both population AMPOP02 and AMPOP10 independently was created with the *VennDiagram* package in R.

Linkage between variants was calculated as the squared Pearson correlation coefficient of the allele frequencies per sample of the variant of interest (*i.e.* the three overlapping variants) with allele frequencies per sample of the other variants on scaffold 2. The threshold of *R^2^* = 0.8 contained the top 1% correlation coefficients.

### Synteny analysis between the D383 and Rookmaker virulence loci

The synteny between the D383 and Rookmaker genomes was computed using ntSynt v1.0.2 (Coombe et al., 2024) with the expected percentage divergence (*-d*) set at 3. All syntenic blocks smaller than 10kb were filtered out and the remaining blocks with synteny to the virulence associated scaffolds (D383 sc2 and Rookmaker sc28) were visualised using *ggplot2* (v3.5.2, https://git.wur.nl/molde006/nemasynt).

### Manual genome annotation of the avirulence locus

Within the 311 kb region of interest on the Rookmaker genome, automated genome annotation predicted 53 transcripts on 48 genes. Because of the inherent limitations of automated genome annotations, the region of interest was manually inspected and curated as described by (Moya et al., 2023).

In the manual curation process, 19 transcripts maintained their Braker predictions, and an additional 23 transcripts had only untranslated regions (UTRs) added. For 5 transcripts, structural changes were applied to one or more exons. Furthermore, 3 transcripts resulted from gene fusion events and 4 transcripts emerged from the splitting of two genes. Notably, 19 extra transcripts were predicted for genes initially identified by Braker, and the manual curation process identified 3 genes not identified by braker. This curation effort yielded a total of 76 transcripts for 47 genes.

### Protein sequence analyses for examining the genes on the avirulence locus

Signal peptide prediction was carried out on the amino acid sequences using SignalP - 6.0 selecting ‘Eukarya’ as organism (Teufel et al., 2022). Transmembrane prediction was carried out using TMHMM - 2.0 on default settings (Krogh et al., 2001). BLASTx was performed with the coding nucleotide sequences against the non-redundant protein sequences (nr) of the NCBI with default parameters (blast.ncbi.nlm.nih.gov/Blast.cgi). BLASTn was performed with the genomic transcript nucleotide sequences against the nucleotide collection (nr/nt) of the NCBI with default parameters (blast.ncbi.nlm.nih.gov/Blast.cgi). Domain predictions were conducted with Interpro (Paysan-Lafosse et al., 2023).

### Temporal transcriptome analysis in avirulent and virulent G. pallida juveniles

To assess the transcriptome of pre-parasitic and parasitic *G. pallida* juveniles *in vitro* infection assays were performed as described before (Zheng et al., 2022). In brief, Rookmaker and AMpop02_DS4_ cysts were hatched based on established protocols (Goverse et al., 2000). Pre-parasitic second stage juveniles (ppJ2s) were collected, cleaned by sucrose purification (Jenkins, 1964) and surface-sterilized in 0.008% (w/v) mercuric chloride. Fourteen-day-old stem cuttings of the potato cultivar Seresta were inoculated with 100 surface-sterilized Rookmaker or AMPOP02 ppJ2s. After inoculation, plants were kept in the dark at 18 °C, and infected root segments were harvested 1-, 3-, 6-, and 9 days post inoculation (dpi). Each sample consisted of the infected root tissue pulled from 10 plants. Four independent hatchings were used to conduct four time-separated biological replicates of the experiment. For each batch, a subsample of the inoculum was used to assess transcription in ppJ2s. Total RNA was extracted with the Maxwell 16 LEV-plant RNA kit (Promega, USA), according to the manufacturer’s instructions. Samples were split, to send ≥300 ng of total RNA for transcriptome sequencing and ≥1µg of total RNA for sRNA sequencing to BGI Genomics (China). For transcriptome sequencing, stranded libraries were sequenced on the DNBseq G400 platform, generating an average of 77.7 million paired-end 150 base pair reads per sample (**Supplementary table 13**). The reads were mapped against the *G. pallida* Rookmaker genome (v.0.8.1), using hisat2 (v2.2.1; Kim et al., 2019). After quality control by MultiQC (Ewels et al., 2016), all samples were used for further analysis.

BAM files were loaded into SeqMonk (v.1.48.1, https://www.bioinformatics.babraham.ac.uk/projects/seqmonk/). Raw count values were generated using the RNA-Seq quantitation pipeline in SeqMonk. Differentially expressed genes were identified by DESeq2 at each of the four parasitic timepoints (Love et al., 2014). P-values were corrected for multiple testing (FDR) and independent filtering was applied. TPM values were extracted from SeqMonk and loaded into R (v.4.3.2; R Core Team, 2013) for data visualisation.

### Silencing of Gp-pat-1 in pre-parasitic juveniles

To identify unique regions in the coding sequences, we performed a BLAST with the coding sequence (CDS) against the *G. pallida* Rookmaker CDS database (21,026 sequences). The siRNA generator of Eurofins Genomics (https://eurofinsgenomics.eu/en/ecom/tools/sirna-design/) was used to design siRNA. The siRNAs are 21 nucleotides long and have an UU dinucleotide at the 3’end (**Supplementary table 9**). Single stranded RNA of the sense and antisense strands were ordered at IDT Europe (Belgium). siRNA targeting eYFP was included as a negative control.

Approximately 5000 juveniles were soaked in 167 µM double stranded siRNA, being either 83.3 µM of each of the effector-targeting siRNA or 167 µM of the eYFP-targeting siRNA. After two hours incubation, nematodes were surface sterilised with mercury chloride for 20 minutes. Approximately, 100 juveniles were inoculated per 14-day-old potato cutting. Approximately 500 juveniles were used for RT-qPCR. Primer sequences for housekeeping genes *Gp_AMA1* and *Gp_GR* were obtained from Sabeh et al. (2018; **Supplementary table 9**). Gene-specific qPCR primers were obtained from IDT Europe (Belgium). Infection assays were performed as described before, with the only change that potato cuttings were grown on Gamborg B5 with 10 g/L sucrose instead of 20 g/L, as this limited the number of avirulent females that were able to develop on resistant potato varieties. Females were counted at 35 dpi.

To assess the expression levels of *Gp-pat-1* at 3 dpi, infected root tissue of 10-15 plants was pooled into one biological replicate. RNA was extracted as described before. We synthesized cDNA using GoScript™ Reverse Transcriptase (Promega, USA), and performed qPCR using the iQ™ SYBR® Green Supermix (Bio-Rad, USA) according to the manufacturer’s protocols. *Gp-pat-1* expression levels were normalized based on the expression of the two housekeeping genes using the 2^−ΔΔC^_T_ method (Livak & Schmittgen, 2001). We normalized the gene expression per batch based on the median expression measured in the eYFP-silenced plants.

## Results

### Commercial potato resistance was overcome by virulent G. pallida field populations

Over the past decade, several European countries have reported the emergence of resistance-breaking *G. pallida* populations, both in the field and under experimental conditions (den Nijs & van Heese, 2019; Fournet et al., 2013; Niere et al., 2014; Phillips & Blok, 2008). To assess virulence levels of *G. pallida* field populations on *G. pallida*-resistant potato varieties, we conducted a first standard PCN resistance test with seven *G. pallida* populations on fifteen Pa2/3-resistant potato varieties and the susceptible variety Desiree (**Figure 1A; Supplementary table 1**). The test panel included two standard populations (Chavornay and E400 ‘Rookmaker’), and five field populations. The three *G. pallida* field populations with suspected virulence (AMPOP02, AMPOP13 and 2017dC) consistently showed higher reproduction rates (P_f_/P_i_) compared to the standard populations (two-sided t-test, p < 0.001; **Supplementary figure 1**). Since virulence can be defined by a population’s reproductive ability on a resistant host relative to its reproductive ability on a susceptible host (Schouten & Beniers, 1997), we concluded that these field populations are virulent on a broad range of potato varieties.

**Figure 1:**
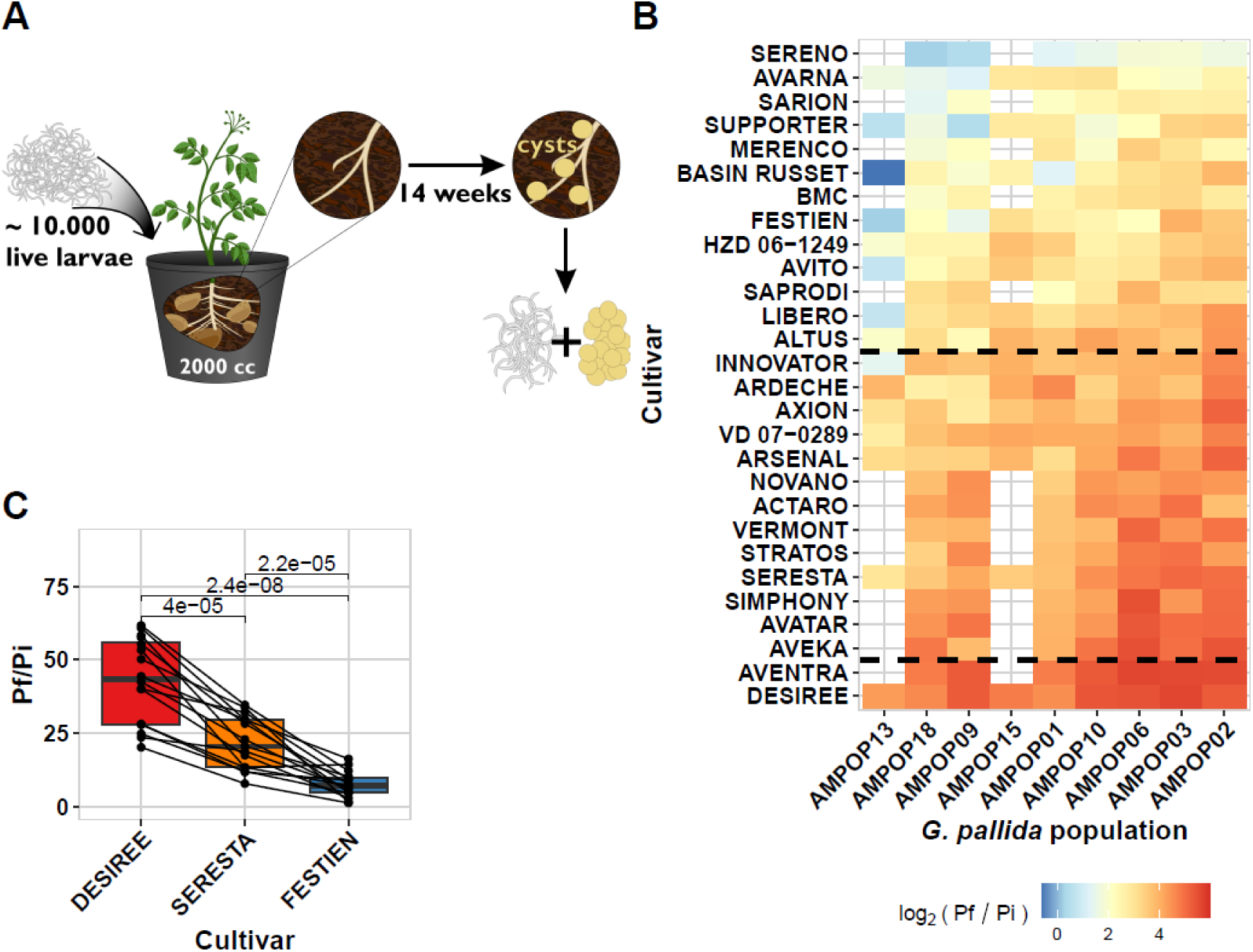
*G. pallida* populations from the Netherlands display virulence on resistant potato varieties. (**A**). The set-up of the standard PCN resistance tests. **(B**) A heatmap of the average log_2_ (Pf/Pi) of the *G. pallida* field populations per potato variety. The blue colours indicate lower propagation and the red colours indicate higher propagation. The dashed lines indicate the separation between the Desiree-, Seresta-, and Festien clusters (**Supplementary figure 3**). Raw data can be found in **Supplementary table 2.** (**C**) The propagation (Pf/Pi) of the virulent *G. pallida* field populations on Desiree, Seresta, and Festien. Each dot represents the mean propagation (three replicates) of a *G. pallida* population within a year (2017 or 2018) on the particular variety. The boxplots summarize the data per variety (15-16 datapoints per variety). The lines connect the measurements per *G. pallida* population within a year. The significances displayed are from a t-test.

We hypothesise that variations in virulence levels can be explained by differences in the genetic backgrounds of the tested potato varieties. To test this, we conducted a second and third standard PCN resistance test in which we tested nine virulent *G. pallida* field populations on a total of 28 potato varieties. Our results showed that the most virulent *G. pallida* populations were able to propagate on all resistant potato varieties (**Figure 1B**; **Supplementary figure 2; Supplementary table 2**). Moreover, virulence on one resistant variety seems to correlate with virulence on other resistant varieties. To assess these correlations, we performed cluster analysis including all 28 potato varieties. This revealed three distinct resistance groups (**Supplementary figure 3**). The groups could be defined by three commonly used varieties for virulence testing: Desiree (no resistance), Seresta (medium resistance), and Festien (strong resistance). Therefore, we named the clusters after these three varieties, where we always saw that the resistance of Festien (Cl_FES_) > Seresta (Cl_SER_) > Desiree (Cl_DES_) (**Figure 1C**; Two-sided t-test, p < 0.001).

Based on these findings, we pursued two lines of further enquiry. First, we investigated the origin of the resistance overcome by virulent *G. pallida* populations. Given our observations, we hypothesized that virulence in *G. pallida* field populations arose from the breakdown of a single resistance shared by all formerly resistant potato varieties. Second, we set out to identify the avirulence locus under selection in *G. pallida*. Given the parallel observation of virulent populations in Germany and The Netherlands (Mwangi et al., 2019), we hypothesised that virulence is the result of selection on standing genetic variation, rather than the emergence of novel mutations at different locations. Hence, we expected to be able to identify a single avirulence locus. In the remainder of this report, we will focus on testing these two hypotheses.

### Virulence in *G. pallida* field populations is based on the breakdown of a single resistance; *GpaV_vrn_*

To determine whether virulence in *G. pallida* field populations arises from the breakdown of a single resistance shared by all potato varieties, we examined the genetic background of the tested potato varieties. Pedigree analysis revealed that all resistant varieties had contributions from various *S. vernei* lines, including LGU 8, 20/24 and I-3 (**Figure 2A**; Van Berloo et al., 2007). This finding aligns with previous research that showed that the *GpaV* resistance from *S. vernei* (*GpaV_vrn_*) was no longer effective against two French *G. pallida* populations selected for virulence on the *GpaV_vrn_*-resistant potato variety Ildhér (Fournet et al., 2013). Similarly, *G. pallida* populations recently isolated in Germany, displayed virulence on the *GpaV_vrn_*-containing Seresta, similar to the populations described here (Mwangi et al., 2019). We confirmed that all resistant varieties are positive for the HC marker, indicating that *GpaV_vrn_* is a major source of *G. pallida* resistance across these varieties (Sattarzadeh et al., 2006).

**Figure 2:**
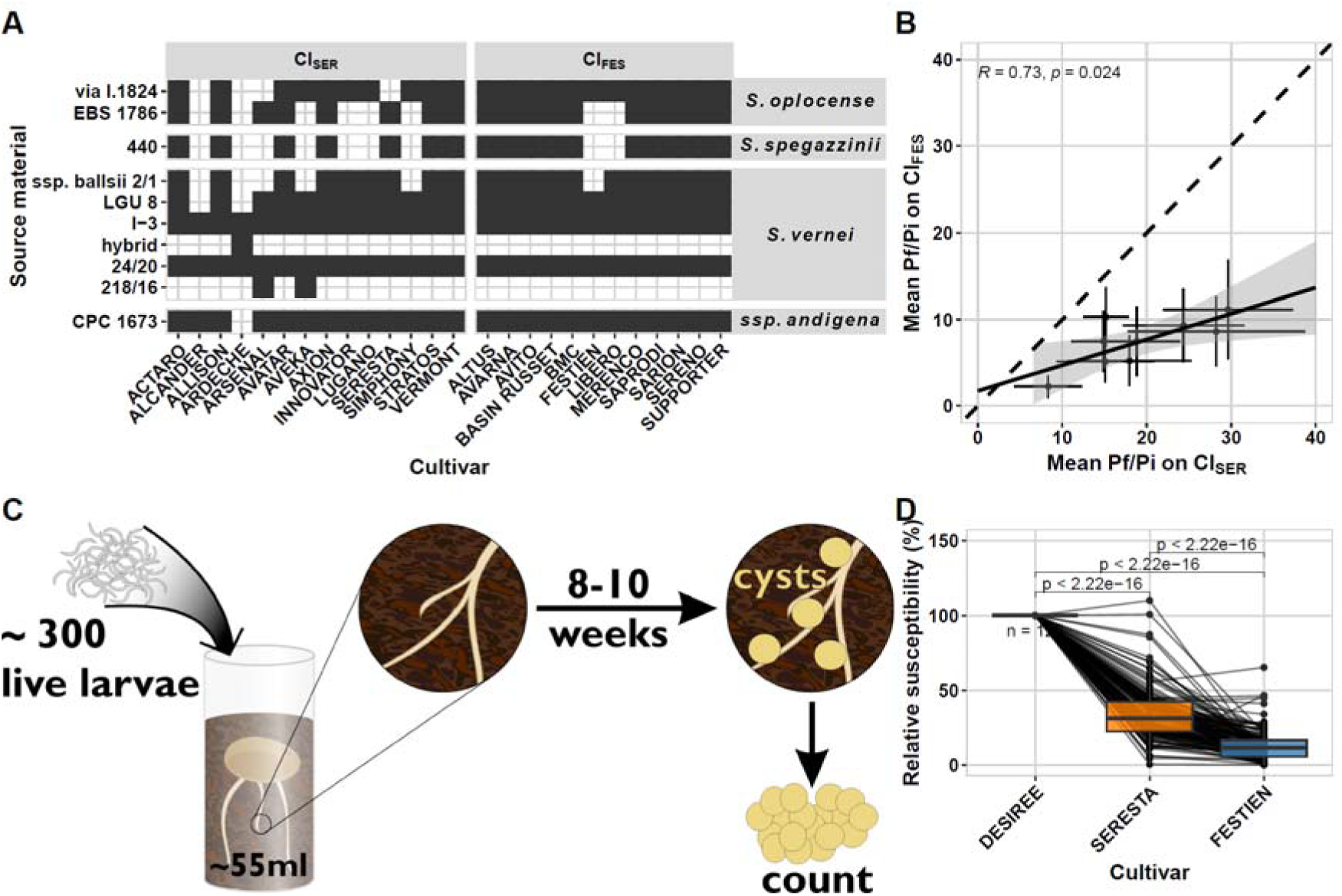
Selection on GpaV_vrn_ can explain virulence in Dutch *G. pallida* field populations. (**A**) The known resistance sources used as progenitors in the tested resistant potato varieties. The data was obtained from the potato pedigree database, which contained pedigrees for 27 of the 31 tested potato varieties (Van Berloo et al., 2007). The potato varieties are separated into the previously identified clusters and the progenitors on the species of origin (ssp. *andigena* is a sub-species of *Solanum tuberosum*). (**B**) A correlation of the mean propagation (Pf/Pi) of nine *G. pallida* populations on the Cl_SER_ varieties with the Cl_FES_ varieties. The coefficient of correlation shown is from a Pearson correlation, visualized in the solid black line. The dashed black line is added as a visual aid. (**C**) The set-up of the small container tests conducted by HLB B.V. to test potentially virulent field populations of *G. pallida*. (**D**) outcome of the small container tests for 125 *G. pallida* populations tested over 2016-2021. Each dot represents the mean relative susceptibility based on 2 – 16 replicates (median of 8 replicates). There were only nine populations where either RS_Seresta_ > RS_Desiree_ or RS_Festien_ > RS_Seresta._ The significances displayed are from a two-sided t-test.

**Figure 3:**
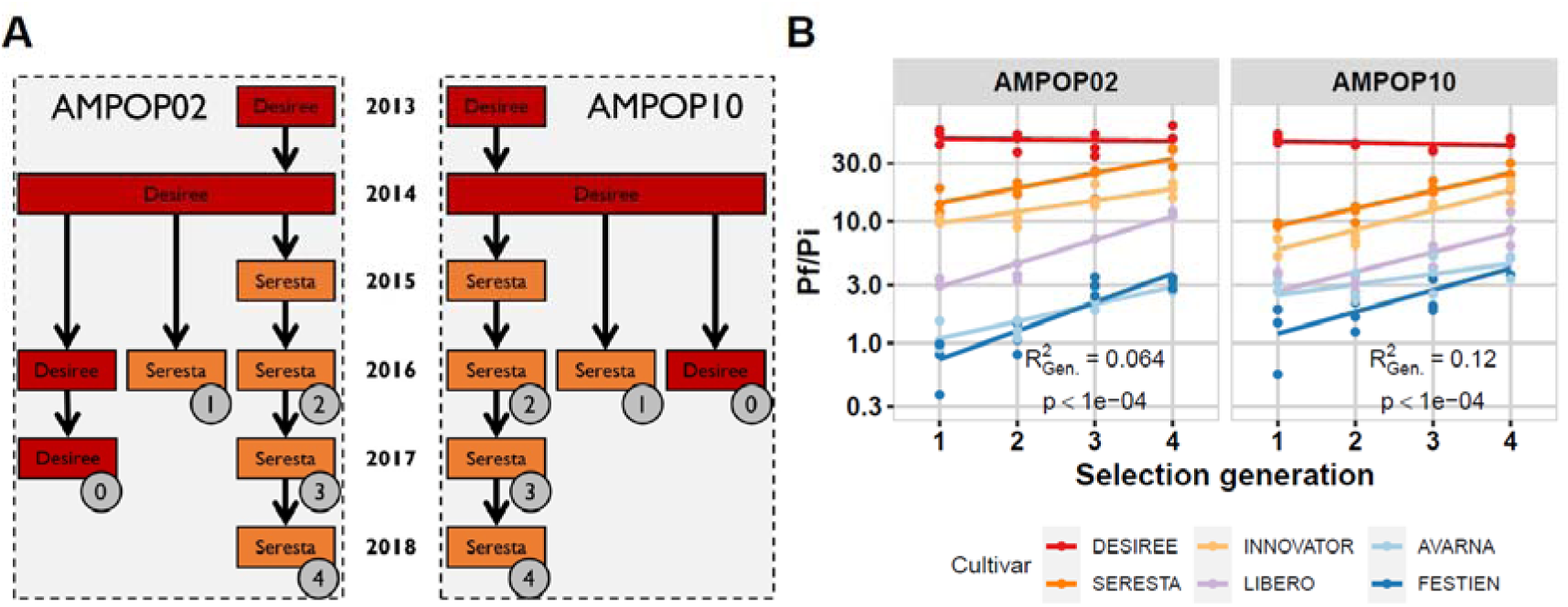
Selection on the *GpaV_vrn_*-containing potato variety Seresta also increases virulence on other commercial potato varieties. (**A**) The selection experiment that was carried out with four generations of selection on Seresta. The colours indicate the potato variety used (Red Desiree, Orange Seresta) and the circles with the numbers indicate which propagation was used in the standard PCN resistance test. The numbers indicate the generation of selection. Due to the amount of material required for the standard PCN resistance test, generation 0 and generation 1 could not be derived from the selection line generations 2-4 were on. (**B**) The outcome of the standard PCN resistance test for selected generations 1-4. Each dot represents the reproduction (Pf/Pi) measured in one two-litre pot inoculated with 10,000 larvae. The lines are linear regressions added as a visual aid. Colours indicate varieties, where Seresta and Innovator belong to Cl_SER_ and Libero, Avarna and Festien belong to Cl_FES_. The R^2^ is the amount of variance explained by generation as derived from a PERMANOVA. The data can be found in **Supplementary table 7**.

To further verify the presence of *GpaV_vrn_*, we deployed graphical mapping (van Eck et al., 2017). We selected positively for Innovator and negatively against Desiree after which 1733 SNPs remained, of which 170 on Chromosome 5. We confirmed that all the resistant cultivars contain alternative alleles around 4.7 - 5.7 Mb on chromosome 5 of the ST4.03 genome, where *GpaV_vrn_* was previously mapped (**Supplementary table 3**; van Eck et al., 2017). As *Gpa6* might also be present, we blasted the CT220 EST (Rouppe Van der Voort et al., 2000) to the ST4.03 genome, which identified a single region from 60925291 - 60925562 Mb on chromosome 9. Given the pedigree, we selected positively for Seresta and negatively against Desiree after which 1729 SNPs remained, of which 140 on chromosome 9. There were some shared SNPs in that area, but it could not be confirmed which cultivars carry *Gpa6* and which do not (**Supplementary table 4**). Taken together, this indicates that resistance is based on *S. vernei*, with *GpaV_vrn_* as the most parsimonious explanation for resistance.

If virulence is the result of selection on different resistances, we would expect virulence on one resistance to be independent of virulence on other resistances. Conversely, if selection against a singular resistance drives virulence, then differences in virulence among nematode populations would be expected to reflect a gradient of selection, representing the increase of single virulence allele. To test this, we correlated the mean virulence measured within Cl_SER_ with the mean virulence measured in Cl_FES_ across *G. pallida* field populations. The virulence levels between the two clusters were strongly correlated (test of correlation, p = 0.024; **Figure 2B**), indicating that virulence on Cl_SER_ varieties and Cl_FES_ varieties involves the same trait. This confirms that the same resistance, *GpaV_vrn_*, has been overcome in all these varieties.

### *G. pallida* virulence is the result of selection on standing variation

We hypothesised that virulence arose through selection on standing variation, defined as genetic variation present in *G. pallida* field populations prior to the introduction of commercial *GpaV_vrn_* resistant varieties on the market thirty years ago. This implies that virulence alleles share the same origin and have spread across different fields over large distances. Alternatively, virulence may have arisen from a *de novo* mutation after the introduction of *GpaV_vrn_* resistant varieties, in which case virulence would be restricted to a limited number of populations confined to a relatively small geographic area. To test this hypothesis, we assayed a total of 125 *G. pallida* field populations, collected between 2016 and 2021 using small container tests (**Figure 2C; Supplementary table 5**). We found that virulence was widespread and followed a similar pattern to that identified by testing nine *G. pallida* populations on 28 potato cultivars (**Figure 2D; Supplementary figure 4**). This indicates that a scenario where selection on standing variation took place is the most likely.

Selection on standing variation also implies that virulence levels in a *G. pallida* population correspond to its progression through the *GpaV_vrn_*-selection process, with each generation leading to increased virulence. We tested this first by cluster analysis on the 9 *G. pallida* populations tested on 28 potato cultivars (**Figure 1B**). This analysis did not yield distinct groups, it instead arranged populations along a gradient from least to most virulent (**Supplementary figure 5**). This indicates that virulence is a quantitative trait shared among many field populations. Next, we tested how much of the variance could be explained by: (i) the averaged virulence of the *G. pallida* population within the Cl_SER_ (**Figure 2B**) and (ii) the assignment of the potato variety to the two clusters (**Supplementary figure 3**). These two factors, along with their interactions, accounted for 45.2% of variance in the data (PERMANOVA, p < 0.0001; **Supplementary table 6**), leaving only 1.3% of the variance over populations unexplained (PERMANOVA, p < 0.0001; **Supplementary table 6**). This suggests that differences between *G. pallida* field populations primarily reflect quantitative variation in their stage of selection of virulence. Collectively, these findings indicate that virulence in *G. pallida* populations is shaped by selection acting on standing genetic variation, present before the introduction of *GpaV_vrn_*-resistant seed potatoes.

### *G. pallida* virulence on Seresta is a selectable, heritable trait conferring virulence on a range of potato cultivars

To verify that populations can be positively selected for virulence and to proceed with mapping of the virulence allele, two field populations (AMPOP02 and AMPOP10) that had shown virulence on *GpaV_vrn_*before (**Figure 1D**), were further selected on Seresta (which contains *GpaV_vrn_* as major resistance locus (Milczarek et al., 2011)). The selection was conducted over the course of five years and yielded populations selected on Seresta for four consecutive generations. Subsets of these populations (where we had sufficient cysts) were tested for virulence in a fourth standard PCN resistance test. For each population, we had three consecutive generations that we could directly compare to determine the increase in virulence (**Figure 3A**). Unfortunately, we could not directly compare the unrelated previous generations, as these had been propagated on the non-resistant Desiree variety.

Selection on Seresta increased the virulence of AMPOP02 and AMPOP10 on both Cl_SER_ and Cl_FES_ varieties (**Figure 3B; Supplementary table 7**). We found that consecutive generations of selection increased the propagation (Pf/Pi) and thereby the relative susceptibility (RS) in multiple potato varieties (Kruskal-Wallis test on Pf/Pi: p < 0.05 for 9 out of 10 tests; on RS: p < 0.05 for 9 / 10 tests; **Supplementary figure 6A** and **B**). The number of eggs per cyst remained stable over the course of selection (Kruskal-Wallis test on Pf/Pi: p > 0.05 for 11 / 12 tests; **Supplementary figure 6C**). Consistent with our previous experiments, selection on Seresta resulted in increased virulence, also on the more resistant Cl_FES_ varieties (PERMANOVA, p < 0.001; **Figure 3B**).

To assess the genetic underpinnings of virulence, we determined the broad sense heritability (*H^2^*) for each potato variety used in the fourth standard PCN resistance tests, the small container tests, and the selection experiment. All analyses indicated a major genetic component to virulence (**Supplementary table 8**). In the selected populations, the *H^2^* was 0.73 and 0.83 for RS on Seresta for AMPOP02 and AMPOP10, respectively (Permutation, q < 0.01). Given that broad-sense heritability is typically high in the case of a single allele, this suggests the selection of a single major virulence allele that explains the heritable variation. This allele, when selected for on Seresta, also increased virulence in other potato varieties, indicating the reliance on a shared source of resistance (*GpaV_vrn_*) as the genetic basis for *G. pallida* resistance in resistant potato varieties.

### Virulence against *GpaV_vrn_* results from selection at a single locus on the *G. pallida* genome

To test whether virulence is the result of selection on a single major virulence allele, we used the selection experiment to detect alternative alleles that were consistently selected for over the generations. To this end, we sequenced *G. pallida* populations collected from the selection experiment. We aimed to sequence all three biological replicates on both Desiree and Seresta separately. This resulted in sequence data for 45 population-variety combinations (**Figure 4A; Supplementary table 9**). We used whole-genome high-coverage sequencing (achieving a median coverage of 162-fold per position per sample) to be able to estimate allele frequencies within our populations. Variants were called using a newly assembled *G. pallida* reference genome, that is based on the Rookmaker (E400) population. The Rookmaker genome is less fragmented and has a higher quality annotation than the previously published D383 genome (van Steenbrugge et al., 2023; **Supplementary table 10**).

**Figure 4:**
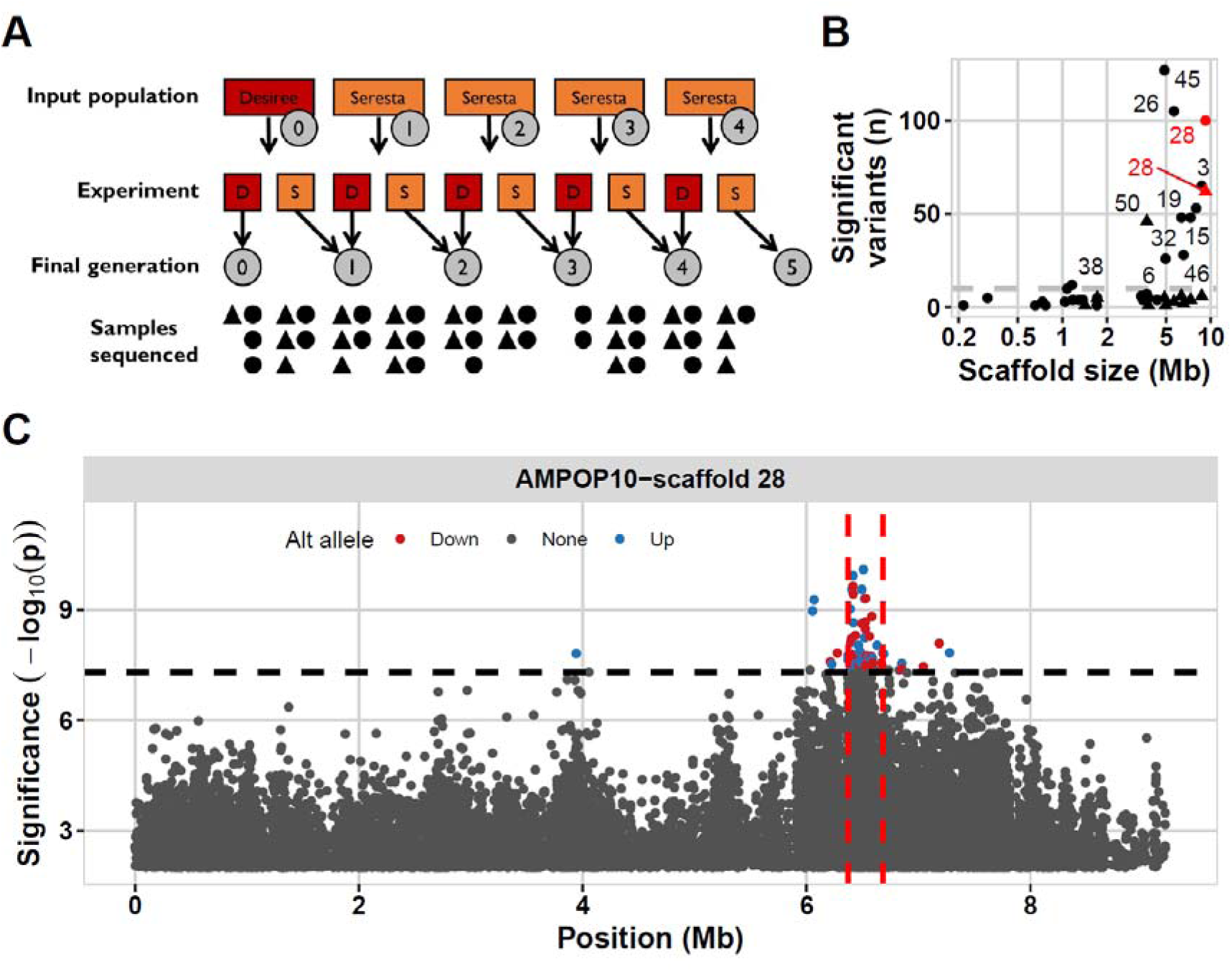
A single genetic locus is selected after repeated selection on Seresta. (**A**) Overview of the AMPOP02 (dots) and AMPOP10 (triangles) samples generated in the selection experiment and how many were successfully sequenced. As the larvae from which we isolated DNA underwent either a selection on Desiree (red) or Seresta (orange), we took the final number of generations selected on Seresta along in the analysis. (**B**) The number of variants that significantly correlate with virulence per scaffold plotted against the scaffold-size in million bases (Mb). Dots indicate the variants found in AMPOP02 and triangles indicate variants were found in AMPOP10. Text indicates which scaffold of the Rookmaker genome the datapoint belongs to. (**C**) The significance of the variants on scaffold 28 associated with generation in AMPOP10. The variants are plotted on their location on scaffold 28 (in Mb) versus the significance in -log_10_(p). The variants where the frequency of the alternative allele is decreasing over generations are coloured red, where the alternative allele is increasing are coloured blue. Grey variants are not significant. The dashed horizontal line indicates the Bonferroni-corrected threshold. The dashed vertical lines indicate the candidate locus. Note that the y-axis has been cut off at -log_10_(p) < 2.

We identified 1,017,636 segregating variants within the two selection populations. We used allele frequencies rather than a standard biallelic matrix to be able to explicitly deal with heterozygous populations – not individuals. We used the allele frequencies in a principal component (PC) analysis to understand the major differences between the 45 samples. We found that the major explanatory component, PC 1 (91.1% of variance) yielded no clear separation and probably captures genetic diversity (heterozygosity) present in both populations (**Supplementary figure 7A**). PC 2 (5.8% of variance) however clearly separated the two populations (AMPOP02 and AMPOP10), probably related to alleles distinct for the two populations. Together PC 3 (0.37% of variance) and PC4 (0.18% of variance) respectively separated the early generations in AMPOP02 and AMPOP10 from the selected populations (**Supplementary figure 7B**). This clearly indicates that there was a genetic signal associated with selection.

To identify alleles associated with virulence we calculated the correlation between alternative allele frequencies and the corresponding generations. For variants associated with virulence, the alternative allele frequencies should increase over the generations of selection as the reference genomes were derived from avirulent *G. pallida* populations. Initially, we identified 680 selected variants in AMPOP02 and 142 selected variants in AMPOP10 (Bonferroni corrected p < 0.05; **Supplementary table 11; Supplementary figure 7C**). In both populations scaffold 28 contained many allelic variants that significantly correlated with virulence (**Figure 4B**). When we investigated the overlap between the significant variants in AMPOP02 and AMPOP10, we found that two variants overlapped, and both were located on scaffold 28 of the Rookmaker genome (hypergeometric test, p < 1*10^-4^; **Supplementary figure 7D-E**). These allelic variants were not located in the coding sequence of a predicted gene, so were not likely to be causal for virulence. However, likely the two variants on scaffold 28 were linked to virulence. We also conducted the association analysis using the *G. pallida* D383 reference genome (van Steenbrugge et al., 2023). Here, the majority of the significantly selected variants were found on a region on scaffold 2 of the D383 genome (**Supplementary figure 8A-E**). We compared the D383 scaffold 2 with the Rookmaker scaffold 28 by assessing the synteny between the two scaffolds (**Supplementary figure 8F**). The identified regions were syntenic, which gives confidence that we identified a single locus linked to *G. pallida* virulence on *GpaV_vrn_*.

We set out to identify the genomic region most likely to contain the virulence allele(s). Therefore, we analysed the combined association in AMPOP02 and AMPOP10 using changepoint analysis. AMPOP10 contained the narrowest introgression of the virulent haplotype (**Figure 4C**), and together with AMPOP02 we found 162 variants on scaffold 28 (**Supplementary figure 9A**). Using these 162 variants, we found the most variant-dense locus between 6.37-6.68 Mb (**Supplementary figure 9B**). Indeed, when we analysed the linkage based on the two overlapping SNPs, multiple variants on Scaffold 28 showed considerable linkage (R^2^ > 0.8; **Supplementary figure 9C-D**). The two overlapping variants on the other hand were not linked (R^2^ = 0.35), which was expected as one was positively- and the other negatively correlated with virulence (**Supplementary figure 7D-E**). Based on these association analyses, we conclude that the search for the causal gene should prioritize the 311 Kb window on scaffold 28 of the *G. pallida* Rookmaker genome.

### Identification of putative resistance-breaking effectors on the virulence-associated locus yields one prime candidate

To identify a virulence allele responsible for breaking resistance, we examined the virulence-associated genomic locus of *G. pallida* for candidate genes. Automated genome annotation predicted 53 transcripts distributed over 48 genes within the 311 Kb locus. Due to the inherent limitations of automated genome annotations, the locus was manually inspected and curated following the approach described by (Moya et al., 2023), resulting in a total of 76 transcripts on 47 genes within the locus (**Supplementary table 12**).

We examined the locus for putative resistance-breaking genes based on three criteria:

(i) their likeliness of being secreted, (ii) their expression during early infection, and (iii) the presence of allelic variation. These criteria align with the understanding that nematodes secrete effector proteins to avoid or supress plant defence responses during early infection, and that virulence results from selection on standing variation.

First, to assess which gene products are possibly secreted, we examined the 76 peptide sequences for the presence of an N-terminal signal peptide for secretion and the absence of transmembrane helices. Based on these criteria, 20 of the 76 transcripts classified as putative effectors (**Figure 5A**; **Supplementary table 12**).

**Figure 5:**
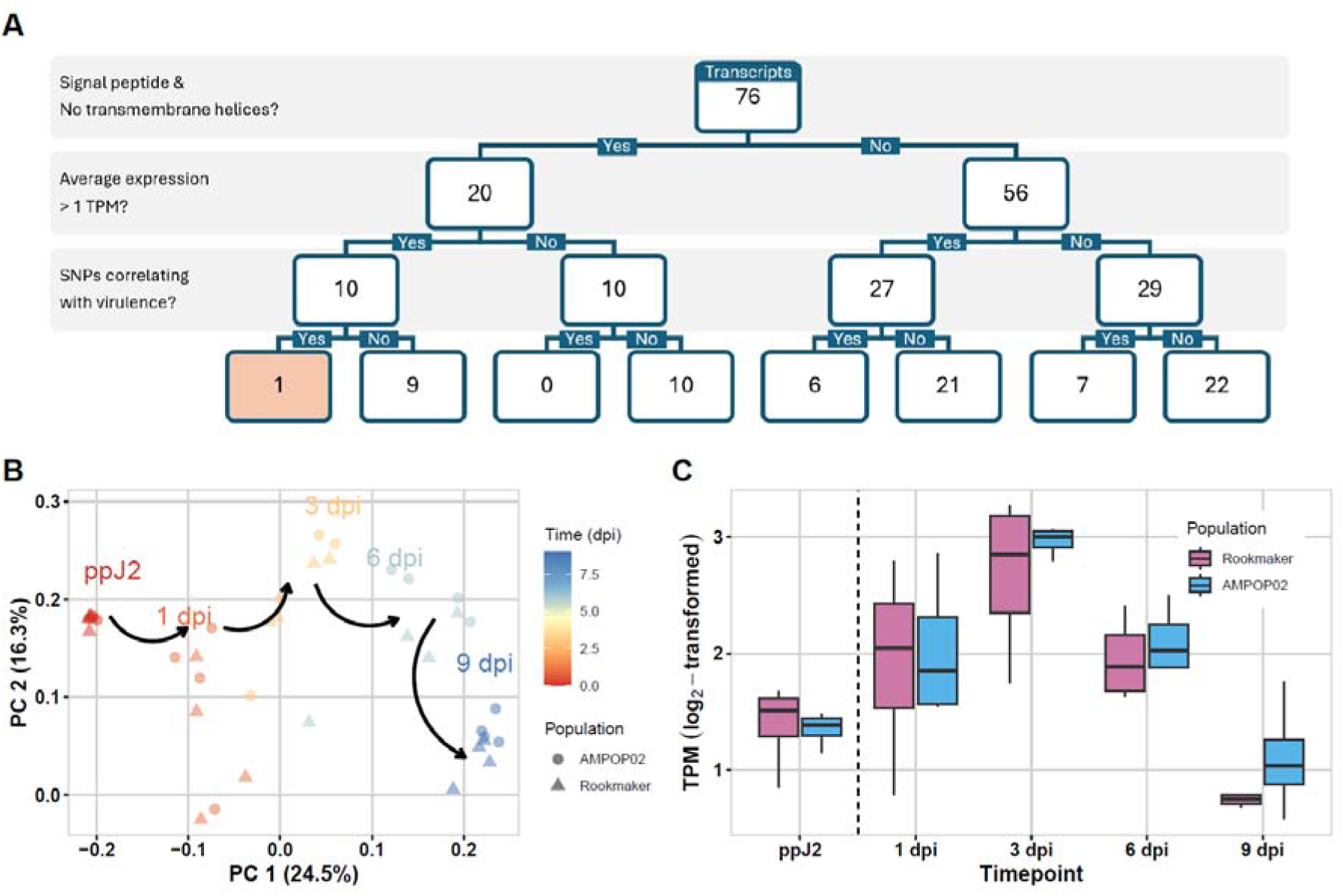
The gene *Gp-pat-1* meets all three criteria of a resistance-breaking effector. (**A**) A flowchart of the three selection criteria leading to the identification of a single candidate (highlighted box). (**B**) Principal component (PC) analysis on RNAseq data of pre-parasitic juveniles (ppJ2) and parasitic juveniles, shows that time captures most of the variance (24.5%). This indicates a transcriptional shift in the nematode as the infection progresses. (**C**) Expression of *Gp-pat-1* in the avirulent Rookmaker and virulent AMPOP02 populations on Seresta. Expression peaks at 3 days post inoculation, roughly coinciding with defence activation of *GpaV_vrn_*.

Second, to assess the expression of the candidate genes during early stages of infection, an infection assay was performed on the *GpaV_vrn_*-resistant potato variety Seresta. Pre-parasitic juveniles (ppJ2s) from avirulent (Rookmaker) and virulent (AMPOP02) *G. pallida* populations were inoculated on two-week old potato cuttings. A subsample of ppJ2s was harvested at inoculation and infected root tissue was harvested at 1-, 3-, 6-, and 9-days post inoculation (dpi) and subjected to transcriptome sequencing. On average, 77.7 million read pairs were generated per sample, with 87.5% of ppJ2 reads and 2.6% of reads from infected root tissue mapping to the *G. pallida* Rookmaker genome (**Supplementary table 13**). Principal Component (PC) analysis revealed that within the dataset time is the factor explaining most of the variance (**Figure 5B**). Interestingly, none of the 76 transcripts showed statistically significant expression differences between Rookmaker and AMPOP02 at any timepoint during infection (FDR<0.01). Given that *S. verneii* resistance has been described as a ‘late’ type of resistance response, activated in the first few days of infection, but not within the first 24 hours (Rice et al., 1987), we hypothesized that the expression of a resistance-breaking effector coincides with defence activation. Therefore, we prioritized candidates with an average log_2_-transformed expression of at least 1.0 transcript per million (TPM) at 3 dpi. Of the 76 transcripts, 37 transcripts passed this threshold (**Supplementary table 14**). Among the 20 putative effectors, 10 transcripts passed the threshold (**Figure 5A**).

Third, we assessed allelic variation and alternative allele frequencies in AMPOP02 and AMPOP10 over the generations in the selection experiment. Since transcriptomic analysis did not indicate differences in expression levels between Rookmaker and AMPOP02, we inferred that virulence likely results from a structurally different effector, rather than a quantitative difference in expression. Therefore we prioritized variants affecting the coding sequences of the genes in the region. Of the 76 transcripts at the locus, 14 contained single nucleotide polymorphisms (SNPs) positively and significantly (FDR<0.01) correlating with virulence in AMPOP02 and AMPOP10 (**Supplementary table 13**). Among the 10 candidates, just 1 contained allelic variation (*Gpal_Rook_g6760.t1-0004* [*Gp-pat-1*]). Interestingly, *Gp-pat-1* also shows the most significant quadratic correlation with time of all putative effectors (Adj. p-value = 9.53e-7; **Figure 5C**), suggesting that expression coincides with the *GpaV_vrn_* defence response. As this is the only gene passing all three selection criteria, this gene is considered the prime candidate for breaking *GpaVvrn* resistance.

### Silencing *Gp-pat-1* enhances virulence on *GpaV_vrn_*-resistant potato varieties

To assess the role of our prime candidate gene in the breakdown of *GpaV_vrn_*-resistance, we silenced *Gp-pat-1* by soaking juveniles of AMPOP02 in small interfering RNAs (siRNAs). Since the *Gp-pat-1* expression in pre-parasitic juveniles is not representative for the expression at 3 dpi (**Figure 5C**), we used infected root tissue to assess expression levels. This resulted in a 0.72-fold expression of our effector at 3 dpi (**Figure 6A**). Although, expression levels were overall lower, there was a lot of biological variation. Upon inoculating Desiree and Seresta, we observed a significant increase of virulence on Seresta, but not on Desiree (**Figure 6B**). The unchanged virulence of *G. pallida* on the susceptible Desiree, combined with increased virulence on the *GpaV_vrn_*-resistant Seresta, indicates that *Gp-pat-1* functions as an avirulence factor recognized by *GpaV_vrn_*.

**Figure 6:**
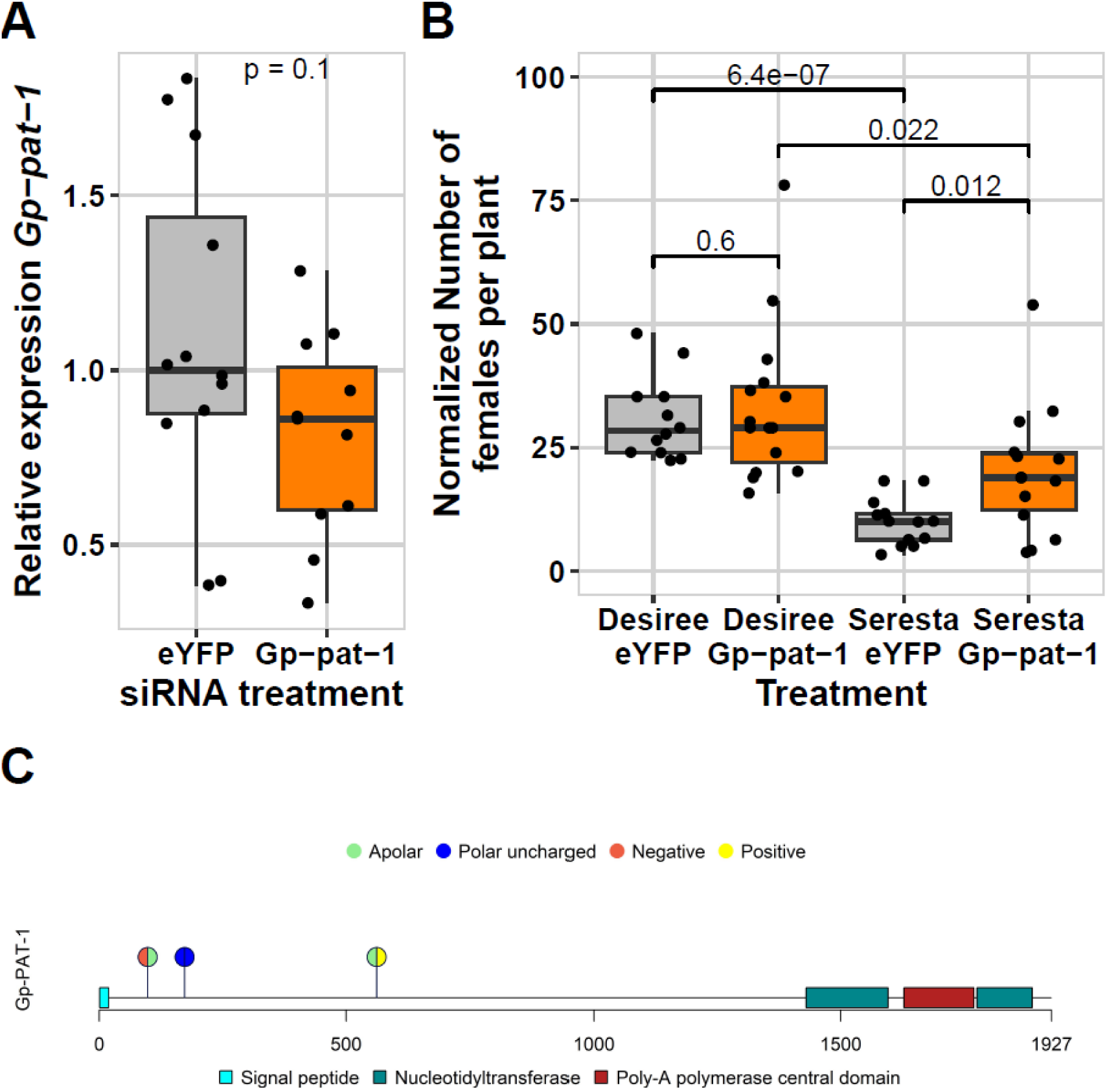
RNAi of *Gp-pat-1* increases virulence on the *GpaV_vrn_*-resistant potato cultivar Seresta. (**A**) Expression levels of *Gp-pat-1* 3 days after inoculation. Expression was relative to eYFP-treated ppJ2. Combined data from two batches is shown. The significance shown is from a t-test. (**B**) Juveniles with an altered *Gp-pat-1* expression were more virulent on the *GpaV_vrn_*-resistant Seresta compared to the control juveniles that are soaked in eYFP targeting siRNAs. Virulence is unchanged on the susceptible cultivar Desiree. The p-values shown are from t-tests. (**C**) Schematic representation of *Gp*-PAT-1 depicting the predicted functional domains and the four nonsynonymous SNPs that correlate significantly with virulence in AMPOP02 and AMPOP10. These four SNPs cause three amino acid changes. Colours of the circles indicate polarity of the amino acid according to the reference sequence (left half) and the alternative sequence (right half).

To further characterize *Gp-pat-1*, we searched for functional domains using Interpro (Paysan-Lafosse et al., 2023). Our effector contains a poly-A polymerase domain, which includes three subdomains: a central poly-A polymerase domain flanked by two nucleotidyltransferase domains on both sides (**Figure 6C**). As the gene has the characteristics of a polynucleotide adenylyltransferase we named it *Gp-pat-1*. The *Gp*-PAT-1 protein is predicted to have nuclear subcellular location, where it may be involved in mRNA 3’ processing.

To identify variants within *Gp-pat-1* potentially causal for virulence, we assessed the allelic variants that significantly correlated with virulence in the selection experiment (**Figure 3; Supplementary table 13**). Within the coding sequence of *Gp-pat-1*, we identified four SNPs leading to non-synonymous changes in the protein. Furthermore, these four SNPs correlated with virulence (FDR<0.01). As two of the SNPs were located within the same codon, these four SNPs caused three amino acid changes, two of which altered the charge of the amino acid (**Figure 6C**). We therefore conclude that *Gp-pat-1* is a strong candidate as *GpaV_vrn_* virulence-associated effector.

## Discussion

### *Globodera pallida* virulence is the result of the widespread use of *GpaV_vrn_*

We show that resistance-breaking *G. pallida* field populations are virulent on a broad range of potato varieties. Moreover, the level of virulence on one resistant variety correlated with the level of virulence on other resistant varieties. This suggests that virulence is the result of the breakdown of a common source of resistance present in all tested potato varieties. Our data identify *GpaV_vrn_*as the resistance broken by virulent *G. pallida*. The broad presence of *GpaV_vrn_* is in line with literature wherein *GpaV_vrn_* was expected to be the primary source of *S. vernei*-derived resistance in French and British *G. pallida*-resistant commercial potato varieties (Fournet et al., 2013; Varypatakis et al., 2019). Other reports have mentioned *Grp1* as the main source of *G. pallida* resistance in commercial potato cultivars (Grenier et al., 2020). *Grp1* co-localizes with *GpaV_vrn_* on chromosome V, but is not present in all varieties tested here and its breakdown is therefore unlikely to explain the increase in virulence observed here (Finkers-Tomczak et al., 2009; van der Voort et al., 1998).

Cluster analysis on the susceptibility levels of 29 potato varieties clustered the potato varieties into two clusters, Cl_SER_ and Cl_FES_, named after the often-grown Seresta and Festien varieties. Resistance levels in Cl_FES_ correlate with – but exceed that of – Cl_SER_, suggesting that Cl_FES_ might have additional minor resistances to *G. pallida* on top of *GpaV_vrn_*. Candidates for this additional resistance include *Gpa6*, *Grp1, GpaV_spl_*, and *GpaXI_spl_* (Caromel et al., 2005; Gartner et al., 2021; Rouppe Van der Voort et al., 2000). Our data point to a scenario of a broad application of *GpaV_vrn_* since the introduction of the first potato varieties carrying *GpaV_vrn_* in 1994 (Van Berloo et al., 2007). This narrow genetic basis of *GpaV_vrn_* resistance, especially in the Cl_SER_, led to the emergence of virulent *G. pallida* field populations. Since adaptation of *G. pallida* populations to Cl_SER_ potato varieties also enhances their virulence on Cl_FES_ varieties, selection on *GpaV_vrn_* not only breaks resistance in CL_SER_ but also increases the potential of these populations to overcome additional resistances present in Cl_FES_ varieties.

### Virulence is widespread and results from selection on standing variation

The origin of virulence alleles has big implications for their geographical distribution. When virulence alleles were present in *G. pallida* field populations prior to the introduction of *GpaV_vrn_*, virulence might be widely distributed, but if virulence alleles arose from a recent *de novo* mutations, virulence likely is restricted to a single field or a limited set of neighbouring fields.

We demonstrate widespread virulence across the Northeast of the Netherlands and that differences in virulence levels reflect differences in stages of selection. Thus, we conclude that virulence is likely the result of selection on standing variation. Virulence is not limited to the Netherlands, the parallel emergence of German resistance-breaking populations (Niere et al., 2014) and the resistance-breaking potential of French and English populations (Fournet et al., 2013; Phillips & Blok, 2008; Varypatakis et al., 2019) indicate that virulence alleles are present in *G. pallida* populations throughout Western Europe.

The spread of virulence across Europe suggests that virulence alleles were already present in the founding populations that were introduced into Europe roughly 150 years ago (Plantard et al., 2008). This is supported by the observation that some Peruvian *G. pallida* populations are virulent on *GpaV_vrn_*, without being exposed to this resistance in Europe (Hockland et al., 2012). Given that all European mainland *G. pallida* populations originated from a single historical introduction (Grenier et al., 2020; Plantard et al., 2008), it is also unlikely that virulence alleles were introduced into Europe in a more recent undescribed introduction of *G. pallida* into Europe. Since all European mainland populations are from the same historic introduction, and the buildup of virulence observed in our selection experiment is comparable to the virulence buildup in other artificial selection experiments with *G. pallida* on *GpaV_vrn_* (Fournet et al., 2013; Varypatakis et al., 2019), *G. pallida* populations. However, to test whether a shared virulence allele is responsible for virulence across Western Europe, future research should compare the genetic basis of virulence in Dutch populations with that of other recent West-European *G. pallida* populations.

### Associating virulence to a single locus suggests a gene-for-gene interaction

Given that virulence results from a single selection pressure in potato, we hypothesised that virulence results from selection on a single *G. pallida* locus. Studies on other nematode species suggest that heritabilities within the range we observed (*H^2^* ≥ 0.73), are often associated with a single genetic locus (Evans et al., 2021). High-coverage whole-genome sequencing on each generation of the two selected populations and the assembly of the new highly contiguous *G. pallida* Rookmaker reference genome allowed us to associate virulence to a single 311 Kb locus. In comparison, two previous genome scans on French *G. pallida* populations virulent on *GpaV_vrn_*, landed on genomic regions of roughly 2 Mb scattered over multiple scaffolds (Eoche-Bosy, Gauthier, et al., 2017; Eoche-Bosy, Gautier, et al., 2017). A third genome scan consisting of two variant calling analyses, PenSeq and ReSeq, resulted in the identification of 58 and 561 allelic variants, respectively, scattered across the genome (Varypatakis et al., 2020).

Linking virulence to a single locus suggests that virulence on *GpaV_vrn_*follows the classical gene-for-gene model (Flor, 1971), in which an effector with avirulence activity triggers a resistance-specific defence response, leading to plant immunity. This model has been widely used for informed resistance deployment and also applies to cyst nematodes (Janssen et al., 1991). In the future, identifying the *GpaV_vrn_* gene from the 9 reported candidates (Wang et al., 2023) will enable testing whether *GpaV_vrn_*and its corresponding avirulence effector conform to the gene-for-gene model.

### The novel effector *Gp-pat-1* has avirulence activity on *GpaV_vrn_*

We identified variation in the *Gp-pat-1* gene as the cause of (a)virulence on *GpaV_vrn_*-resistant potato plants. Our finding builds on the assumption that stylet-secreted effectors of *G. pallida* are the most likely candidates underlying avirulence (Mitchum et al., 2013). Therefore, we only focused on genes encoding proteins containing a predicted signal peptide and lack a transmembrane helix. Furthermore, we based our candidate gene search on expression levels and the presence of genetic variation. Here, it proved to be crucial to work with accurate, manually curated gene models (Moya et al., 2023). The expression of *Gp-pat-1* upon host infestation aligns with typical effector expression patterns, where infective juveniles utilize effectors to migrate through the root, to establish the syncytium, and to suppress host immune responses (Chen & Aravin, 2023; Da Rocha et al., 2021; Gardner, Verma, & Mitchum, 2015; Wen et al., 2021).

To assess avirulence activity associated with *Gp-pat-1* on *GpaV_vrn_*resistant potato varieties, we silenced the gene in pre-parasitic second stage juveniles of *G. pallida* AMPOP02. Silencing of *Gp-pat-1* increased the number of juveniles that developed into mature females on Seresta, which contains *GpaV_vrn_*, but not on the susceptible Desiree. This indicates that the gene has avirulence activity on *GpaV_vrn_* and suggests that either the protein itself or its effects are recognised by *GpaV_vrn_*. Future research can evaluate this hypothesis by *Agrobacterium*-mediated transient expression of different *Gp-pat-1* alleles in a *GpaV_vrn_*background. We did not observe a fitness cost upon silencing *Gp-pat-1* in a susceptible background. This could be due to a biological factor, such as genetic redundancy, or a technical limitation, such as insufficient resolution or statistical power.

Although the role of *Gp*-PAT-1 in the host was beyond the scope of this study, *in silico* analysis provided first insights into protein functioning. Domain prediction identified the domains of a poly-adenylyl transferase. Although no nematode effectors are known to directly affect polyadenylation, as incorrect polyadenylation affects translation, this effector may be secreted to interfere with transcriptional regulation of the host.

## Conclusion

We show that selection on *GpaV_vrn_*-mediated resistance is responsible for the current outbreak of virulence in *G. pallida* field populations in The Netherlands. By associating virulence to a single locus in *G. pallida* and identifying a single gene involved in overcoming *GpaV_vrn_*resistance, our findings open avenues to develop molecular diagnostic tools to monitor *G. pallida* virulence in the field. Our data suggested that selection by *GpaV_vrn_* on standing genetic variation led to the emergence of virulence in *G. pallida*. However, testing this hypothesis requires molecular investigations on historic populations and multiple virulent populations. If we can identify a common molecular basis for virulence, monitoring allele frequencies will become an important agronomical instrument. To ultimately control these virulent *G. pallida* populations, novel resistance genes must be introduced into commercial potato varieties and used strategically.

## Author contributions

ASS, MGS and GS designed the experiments. CCvS and SvdE assisted in DNA isolation. PH conducted the selection experiments. JMvS, DtM, SJSvdR, ASS and MGS conducted the genetic analyses and assembled and annotated the Rookmaker genome. *In vitro* infection assays for RNAseq and functional validation were conducted by ASS. ASS, MGS and GS wrote the paper with input from all other co-authors.

## Acknowledgements

The authors want to thank the private partners in the PALLIFIT project for their support and their constructive attitude towards this project. We want to thank the Branch Organisatie Akkerbouw for funding the third standard PCN resistance test. We want to thank the NVWA for funding the first and fourth standard PCN resistance test. We want to thank Anne Morbach (Schlaugemacht.net) for making Figure 1A and Figure 1E. We want to thank Laura Zuidema, Vera Putker, Amalia Diaz Granados, Hein Overmars, and André Bertran for generating RNAseq data of the E400 Rookmaker population. We also want to thank Timon van Leeuwen, German Rudakov, Vera Putker, Martijn Holterman, Hans Helder, and Aska Goverse for discussions related to this paper.

## Data availability

All scripts and underlying datasets are available through gitlab (https://git.wur.nl/published_papers/Schaveling_2025_Pallifit_virulence). The data of presented experiments has been included in supplementary files. The DNA sequencing data of the selection experiment is deposited at BioStudies (E-MTAB-15408). The Rookmaker genome, the short- and long-read Rookmaker DNA sequencing data (ERR15233941 and ERR15205749, respectively), the structural annotation and the RNA sequencing data used for structurally annotating the Rookmaker genome (ERR15277786) were deposited at the ENA with accession number PRJEB91928. RNAseq of the infection assay presented in this paper was deposited at BioStudies (E-MTAB-15312).

## Supplementary material

**Supplementary figure 1**: The relative susceptibilities of seven *G. pallida* populations on 16 potato varieties as measured in the first standard PCN resistance test. Three field populations with suspected virulence (2017dC, AMPOP13, and AMPOP02) and two field populations without suspicion (2017Te and 2017Pa) were tested versus the standard population ‘Chavornay’. The Dutch standard E400 ‘Rookmaker’ is also included. The significances shown are from a two-sided t-test versus ‘Chavornay’.

**Supplementary figure 2**: The relative susceptibility and propagation of nine virulent *G. pallida* field populations on 28 potato varieties. (**A**) A boxplot of the relative susceptibility of the field populations per potato variety as determined in a second and third standard PCN resistance tests. Each dot represents a replicate experiment. The colours of the boxplots indicate to which cluster the potato varieties belong (**Supplementary figure 3**), red for Cl_DES_, orange for Cl_SER_, and blue for Cl_FES_. The p-values shown are from a kruskall-wallice test. When significant, the tests indicate there is between-population variation for the relative susceptibility. The *G. pallida* populations are ordered based on the clustering analysis from least to most virulent (**Supplementary figure 4**). (**B**) The propagation of the nine *G. pallida* populations on the 28 potato varieties. Colours and significance test as in (**A**). When significant, the tests indicate there is between-variety variation in propagation.

**Supplementary figure 3**: Three clusters of resistant potato varieties explain most of the variance in propagation of virulent *G. pallida* populations. (**A**) Euclidean clustering of the potato varieties based on the propagation levels of nine *G. pallida* field populations on 28 potato varieties. The data was obtained from the second (indicated with Oct_2017) and third (indicated with Oct_2018) standard PCN resistance tests. The clustering reveals three major clusters, ordered on the amount of propagation observed. The highest propagation is observed on non-resistant Desiree and Aventra (Cl_DES_), subsequently there is a cluster containing Seresta (Cl_SER_) and a cluster containing Festien (Cl_FES_). Note that the placement of the Cl_FES_ in the middle is arbitrary. (**B**) The side-by-side differences between the varieties in relative susceptibility were tested using a TukeyHSD test, which was corrected for multiple testing. The numbers indicate the difference in relative susceptibility between the variety on the x-axis with the variety on the y-axis. For instance, the relative susceptibility in Aventra is 23% lower than Desiree. The colours indicate the -log_10_(p) significance, where grey indicates there was no significant difference.

**Supplementary figure 4**: Small container test data shows strong correlation with the data from the second and third standard PCN resistance tests. The relative susceptibilities obtained from the nine virulent *G. pallida* field populations in the standard PCN resistance test were correlated with relative susceptibilities obtained from small container tests. The number of replicates for the standard PCN resistance test was n=3, for the small container tests the median was 8 replicates (between 7 and 14). Each dot represents the mean values for a variety. The colours indicate whether the cultivar belonged to the CL_DES_ (red), Cl_SER_ (orange), or CL_FES_ (blue). Desiree, Seresta, and Festien are coloured in bright red, bright orange, and bright blue, respectively. Two Pearson correlations were calculated, one including Desiree (on top, solid black line) and one excluding Desiree (second line, solid grey line). The dashed black line is added as a visual aid (equal relative susceptibilities from both tests).

**Supplementary figure 5**: Variance in virulence between *G. pallida* populations follows a gradient without clear clustering (**A**) Euclidean clustering of the virulent *G. pallida* populations based on the propagation levels of nine *G. pallida* field populations on 28 potato varieties. The clustering reveals a gradient of virulence, from most virulent to least virulent. Note that the ordering of populations is arbitrary. (**B**) The side-by-side differences between the *G. pallida* field populations in propagation were tested using a Tukey HSD test, which was corrected for multiple testing. The numbers indicate the difference in propagation between the *G. pallida* field population on the x-axis with the *G. pallida* field population on the y-axis. For instance, the AMPOP02 has a Pf/Pi ratio that is on average 15 higher than that of AMPOP13. The colours indicate the -log_10_(p) significance, where grey indicates there was no significant difference.

**Supplementary figure 6**: The reproductive properties of the two *G. pallida* selection populations on six potato varieties. (**A**) The propagation (Pf/Pi) per generation split out for AMPOP02 and AMPOP10. The dashed vertical line separates the three directly descendant generations from the other two generations. The significances are calculated by a Kruskal-Wallis Rank Sum Test. If significant, there are differences between generations. (**B**) As in (**A**) but for the relative susceptibility. (**C**) as in (**A**) but for the number of eggs per cyst.

**Supplementary figure 7**: Identification of the Seresta-selected loci in AMPOP02 and AMPOP10 based on the Rookmaker genome. (**A**) Principal component analysis on the allele frequencies in the sequenced samples of the selection experiment. The first and second principal components (PC) were associated with population used for the selection and together captured 96.9% of variance in the data. Colours indicate generations, samples from AMPOP02 are indicated with a triangle, samples from AMPOP10 with a circle. (**B**) Principal component analysis on the allele frequencies in the sequenced samples. The third and fourth principal components (PC) were associated with generation of selection and together captured 0.73% of variance in the data. Suggesting only a narrow locus was selected. Colours indicate generations, samples from AMPOP02 are indicated with a triangle, samples from AMPOP10 with a circle. (**C**) The overlap in significant variants from the linear model over the generations in AMPOP02 and AMPOP10, split out for increases in Alternative and Reference alleles. Variants derived from AMPOP02 (680) are coloured blue, variants derived from AMPOP10 (142) are coloured green. (**D**) The association of allele frequency of the G nucleotide at position 6518403 of scaffold 28 with relative susceptibility as measured in the fourth resistance pot test. Each dot represents a sample of AMPOP02 and each triangle a sample of AMPOP10. The overall correlation is shown (black dashed line) as well as the correlation of the AMPOP02 (green solid line) and AMPOP10 (blue solid line). Also, the correlation coefficients and significances are given. (**E**) As in (**D**) but for the allele frequency of the A nucleotide at position 6587857 of scaffold 28.

**Supplementary figure 8**: A region associated with virulence on GpaV_vrn_, syntenic to Rookmaker Scaffold 28 was identified on the D383 genome. (**A**) The overlap in significant variants from the linear model over the generations in AMPOP02 and AMPOP10, split out for increases in Alternative and Reference alleles. Variants derived from AMPOP02 (822) are coloured blue, variants derived from AMPOP10 (138) are coloured green. (**B**) The number of variants identified per scaffold, with the scaffold-size on the x-axis (in million bases; Mb) and the number of significant variants on the y-axis. The horizontal dashed grey line indicates 10 variants per scaffold. Circles indicate the variants were found in the AMPOP02 population and triangles indicate variants were found in the AMPOP10 population. Text indicates which scaffold of the D383 genome the datapoint belongs to. (**C**) Identification of the locus most likely to harbour the causal virulence gene(s) based on the significantly associated variants in AMPOP02 and AMPOP10. The x-axis indicates the physical position in million bases (Mb) and the y-axis the order-number (1 - 162). The dashed vertical lines indicate the candidate locus as determined by changepoint-analysis. (**D**) The significance of the variants on scaffold 2 associated with generation in AMPOP02. The variants are plotted on their location on scaffold 2 (in Mb) versus the significance in -log_10_(p). The variants where the frequency of the alternative allele is decreasing over generations are coloured red, where the alternative allele is increasing are coloured blue. Grey variants are not significant. Note that the y-axis has been cut off at -log_10_(p) < 2. (**E**) As in (**D**), but for the variants associated with generation in AMPOP10. **(F**) A synteny plot of scaffold 2 of the D383 genome and scaffold 28 of the Rookmaker genome, with the virulence loci indicated by pink boxes. The annotations indicate which scaffolds are shown (e.g. d2 is scaffold 2 of the D383 genome). Although the genomic context of the two virulence loci differs between the two genomes, both association analyses landed on syntenic regions of the genome. The x-axis indicates the scale in million bases (M).

**Supplementary figure 9**: Region on scaffold 28 associated with virulence. (**A**) The significance of the variants on scaffold 28 associated with generation in AMPOP02. The variants are plotted on their location on scaffold 28 (in Mb) versus the significance in - log_10_(p). The variants where the frequency of the alternative allele is decreasing over generations are coloured red, where the alternative allele is increasing are coloured blue. Grey variants are not significant. The dashed horizontal line indicates the Bonferroni-corrected threshold. The dashed vertical lines indicate the candidate locus. Note that the y-axis has been cut off at -log_10_(p) < 2. (**B**) Identification of the locus most likely to harbour the causal virulence gene(s) based on the significantly associated variants in AMPOP02 and AMPOP10. The x-axis indicates the physical position in million bases (Mb) and the y-axis the order-number (1 - 162). The dashed vertical lines indicate the candidate locus as determined by changepoint-analysis. (**C**) Linkage between the G nucleotide at position 6518403 of scaffold 28 and the variants on scaffold 28. The x-axis shows the position of the variants in million bases (Mb). The y-axis shows the squared Pearson correlation. The y-axis was cut-off at R^2^ < 0.4. The dashed horizontal line indicates the threshold of the 1% best-correlating variants (R^2^ = 0.56). The dashed vertical lines indicate the area where the virulence gene is likely to be located as determined by changepoint analysis. The two virulence-associated variants are coloured red. Note that the two have a low correlation (R^2^ = 0.35). (**D**) As in (**C**) but for the A nucleotide at position 6587857 of scaffold 28.

**Supplementary table 1**: Data of the first resistance pot test. In total 7 *G. pallida* populations were tested on 16 potato cultivars.

**Supplementary table 2**: Data of the second and third resistance pot tests. In total 28 potato cultivars were tested on 9 *G. pallida* populations.

**Supplementary table 3**: The variants linked to the *GpaV_vrn_* locus as identified by graphical mapping using Desiree and Innovator as references. The positions of the array probes on the ST4.03 genome are indicated, as well as the allele dosages for the 22 tested cultivars. Furthermore, previously identified *GpaV_vrn_*-linked SNPs are indicated (van Eck et al., 2017). The linked presence of alternative alleles makes it likely that all 21 potato cultivars carry the *GpaV_vrn_*resistance.

**Supplementary table 4**: The variants linked to the *Gpa6* locus as identified by graphical mapping using Desiree and Seresta as references. The positions of the array probes on the ST4.03 genome are indicated, as well as the allele dosages for the 22 tested cultivars. The *Gpa6* locus maps to the end of chromosome 9. It is not clear which cultivars carry do - or do not - carry *Gpa6*.

**Supplementary table 5**: Data of the small container tests executed by HLB B.V. The dataset contains data of 125 *G. pallida* populations on the cultivars Desiree, Seresta, and Festien.

**Supplementary table 6**: Analysis of variance for potato variety cluster and *G. pallida* field population virulence. The outcome of a PERMANOVA analysis, showing the variance explained per term in the model. The P-values were derived from 10,000 permutations.

**Supplementary table 7**: Data of the fourth resistance pot tests. Two *G. pallida* selection panels (AMPOP02 and AMPOP10), after various generations of selection were tested on six potato cultivars.

**Supplementary table 8**: Heritability analyses on the fourth standard PCN resistance test data, selection experiment populations and the small container test data. The broad-sense heritability estimate is given for each of the six potato varieties. The value between brackets denotes the q-value (false discovery rate) of the heritability. Values that were not significant were shown in red.

**Supplementary table 9**: An overview of the DNA sequenced samples of the selection experiment. The associated sequence files are listed, the population the sequencing belongs to (Population), the variety the population was grown on, the total number of reads generated (in millions), the total number of reads that mapped (in millions), the median coverage on the D383 genome, and comments.

**Supplementary table 10:** Comparative genome statistics of the Rookmaker genome assembly from the current paper and the D383 genome (van Steenbrugge et al., 2023).

**Supplementary table 11:** The significantly associated variants with increases and decreases in alternative allele frequencies over generations. The table lists the selection population the variant was found in (either AMPOP02 or AMPOP10), the scaffold number, the position of the variant and the reference nucleotide (D383) and the alternative nucleotide (found in the selection population). The output of the linear model over the generations is also given, the significance (uncorrected), the increase in alternative allele frequency (fraction per generation) and the Bonferroni-corrected significance.

**Supplementary table 12:** An overview of the 76 manually annotated transcripts on the avirulence locus, including their position, method of annotation, how the transcripts scored for each of the selection criteria and the best informative BLASTp hit.

**Supplementary table 13:** An overview of the RNA samples used for temporal transcriptome analysis, including sequencing and mapping statistics. All RNAseq data was mapped to the Rookmaker genome and the DM1-3 genome (v4.03; Xu et al., 2011).

**Supplementary table 14:** The average log_2_ transformed transcription of each transcript at each of the five time points measured (i.e., pre-parasitic, 1 day post inoculation (dpi), 3 dpi, 6 dpi, and 9 dpi) for both the avirulent Rookmaker population and the virulent AMPOP02 population. Candidate genes for virulence on *GpaV_vrn_* were required to have a minimum transcription of at least 1 transcript per million (TPM).

**Supplementary table 15:** An overview of the RNA oligonucleotides used for silencing *Gp- pat-1* and DNA oligonucleotides used for RT-qPCR.

## References

Alonge, M., Lebeigle, L., Kirsche, M., Jenike, K., Ou, S., Aganezov, S., Wang, X., Lippman, Z. B., Schatz, M. C., & Soyk, S. (2022). Automated assembly scaffolding using RagTag elevates a new tomato system for high-throughput genome editing. Genome biology, 23(1), 258. 10.1186/s13059-022-02823-7

Bates, D., Mächler, M., Bolker, B., & Walker, S. (2015). Fitting linear mixed-effects models using lme4. Journal of statistical software, 67, 1–48. 10.18637/jss.v067.i01

Brůna, T., Hoff, K. J., Lomsadze, A., Stanke, M., & Borodovsky, M. (2021). BRAKER2: automatic eukaryotic genome annotation with GeneMark-EP+ and AUGUSTUS supported by a protein database. NAR genomics and bioinformatics, 3(1). 10.1093/nargab/lqaa108

Caromel, B., Mugniéry, D., Kerlan, M.-C., Andrzejewski, S., Palloix, A., Ellissèche, D., Rousselle-Bourgeois, F., & Lefebvre, V. (2005). Resistance Quantitative Trait Loci Originating from Solanum sparsipilum Act Independently on the Sex Ratio of Globodera pallida and Together for Developing a Necrotic Reaction. Molecular Plant-Microbe Interactions®, 18(11), 1186–1194. 10.1094/mpmi-18-1186

Coombe, L., Kazemi, P., Wong, J., Birol, I., & Warren, R. L. (2024). Multi-genome synteny detection using minimizer graph mappings. bioRxiv, 2024.2002.2007.579356. 10.1101/2024.02.07.579356

Coombe, L., Li, J. X., Lo, T., Wong, J., Nikolic, V., Warren, R. L., & Birol, I. (2021). LongStitch: high-quality genome assembly correction and scaffolding using long reads. BMC bioinformatics, 22(1), 534. 10.1186/s12859-021-04451-7

Cotton, J. A., Lilley, C. J., Jones, L. M., Kikuchi, T., Reid, A. J., Thorpe, P., Tsai, I. J., Beasley, H., Blok, V., & Cock, P. J. (2014). The genome and life-stage specific transcriptomes of Globodera pallidaelucidate key aspects of plant parasitism by a cyst nematode. Genome biology, 15(3), 1–17. 10.1186/gb-2014-15-3-r43

Danecek, P., Auton, A., Abecasis, G., Albers, C. A., Banks, E., DePristo, M. A., Handsaker, R. E., Lunter, G., Marth, G. T., & Sherry, S. T. (2011). The variant call format and VCFtools. Bioinformatics, 27(15), 2156–2158. 10.1093/bioinformatics/btr330

Danecek, P., Bonfield, J. K., Liddle, J., Marshall, J., Ohan, V., Pollard, M. O., Whitwham, A., Keane, T., McCarthy, S. A., & Davies, R. M. (2021). Twelve years of SAMtools and BCFtools. Gigascience, 10(2). 10.1093/gigascience/giab008

den Nijs, L., & van Heese, E. (2019). Development of new virulent populations in the Netherlands: how to detect this phenomenon in potato cyst nematodes (PCN)?

Eoche-Bosy, D., Gauthier, J., Juhel, A. S., Esquibet, M., Fournet, S., Grenier, E., & Montarry, J. (2017). Experimentally evolved populations of the potato cyst nematode Globodera pallida allow the targeting of genomic footprints of selection due to host adaptation. Plant Pathology, 66(6), 1022–1030. 10.1111/ppa.12646

Eoche-Bosy, D., Gautier, M., Esquibet, M., Legeai, F., Bretaudeau, A., Bouchez, O., Fournet, S., Grenier, E., & Montarry, J. (2017). Genome scans on experimentally evolved populations reveal candidate regions for adaptation to plant resistance in the potato cyst nematode Globodera pallida. Molecular Ecology, 26(18), 4700–4711. 10.1111/mec.14240

EPPO. (2013). PM 7/40 (3) Globodera rostochiensis and Globodera pallida. EPPO Bulletin, 43(1), 119–138. 10.1111/epp.12025

Evans, K. S., van Wijk, M. H., McGrath, P. T., Andersen, E. C., & Sterken, M. G. (2021). From QTL to gene: <em>C. elegans</em> facilitates discoveries of the genetic mechanisms underlying natural variation. Trends in Genetics, 37(10), 933–947. 10.1016/j.tig.2021.06.005

Ewels, P., Magnusson, M., Lundin, S., & Käller, M. (2016). MultiQC: summarize analysis results for multiple tools and samples in a single report. Bioinformatics, 32(19), 3047–3048. 10.1093/bioinformatics/btw354

Finkers-Tomczak, A., Danan, S., van Dijk, T., Beyene, A., Bouwman, L., Overmars, H., van Eck, H., Goverse, A., Bakker, J., & Bakker, E. (2009). A high-resolution map of the Grp1 locus on chromosome V of potato harbouring broad-spectrum resistance to the cyst nematode species Globodera pallida and Globodera rostochiensis. Theoretical and Applied Genetics, 119(1), 165–173. 10.1007/s00122-009-1026-1

Flor, H. (1971). Current status of the gene-for-gene concept. Annu. Rev. Phytopathol, 9(1), 275–296. 10.1146/annurev.py.09.090171.001423

Fournet, S., Eoche-Bosy, D., Kerlan, M.-C., Grenier, E., & Montarry, J. (2018). Phenotypic and Genomic Modifications Associated with Globodera pallida Adaptation to Potato Resistances. Potato Research, 61(1), 65–71. 10.1007/s11540-018-9358-3

Fournet, S., Kerlan, M.-C., Renault, L., Dantec, J.-P., Rouaux, C., & Montarry, J. (2013). Selection of nematodes by resistant plants has implications for local adaptation and cross virulence. Plant Pathology, 62(1), 184–193. 10.1111/j.1365-3059.2012.02617.x

Gartner, U., Armstrong, M. R., Sharma, S. K., Jones, J. T., Blok, V. C., Hein, I., & Bryan, G. J. (2024). Characterisation and mapping of a Globodera pallida resistance derived from the wild potato species Solanum spegazzinii. Theoretical and Applied Genetics, 137(5), 106. 10.1007/s00122-024-04605-0

Gartner, U., Hein, I., Brown, L. H., Chen, X., Mantelin, S., Sharma, S. K., Dandurand, L.-M., Kuhl, J. C., Jones, J. T., & Bryan, G. J. (2021). Resisting potato cyst nematodes with resistance. Frontiers in Plant Science, 12, 483. 10.3389/fpls.2021.661194

Goverse, A., Overmars, H., Engelbertink, J., Schots, A., Bakker, J., & Helder, J. (2000). Both induction and morphogenesis of cyst nematode feeding cells are mediated by auxin. Molecular Plant-Microbe Interactions, 13(10), 1121–1129. 10.1094/mpmi.2000.13.10.1121

Goverse, A., & Smant, G. (2014). The activation and suppression of plant innate immunity by parasitic nematodes. Annual Review of Phytopathology, 52, 243–265. 10.1146/annurev-phyto-102313-050118

Grenier, E., Kiewnick, S., Smant, G., Fournet, S., Montarry, J., Holterman, M., Helder, J., & Goverse, A. (2020). Monitoring and tackling genetic selection in the potato cyst nematode Globodera pallida. EFSA Supporting Publications, 17(6). 10.2903/sp.efsa.2020.EN-1874

Hockland, S., Niere, B., Grenier, E., Blok, V., Phillips, M., den Nijs, L., Anthoine, G., Pickup, J., & Viaene, N. (2012). An evaluation of the implications of virulence in non-European populations of Globodera pallida and G. rostochiensis for potato cultivation in Europe. Nematology, 14(1), 1–13. 10.1163/138855411X587112

Holterman, M., van der Wurff, A., van den Elsen, S., van Megen, H., Bongers, T., Holovachov, O., Bakker, J., & Helder, J. (2006). Phylum-Wide Analysis of SSU rDNA Reveals Deep Phylogenetic Relationships among Nematodes and Accelerated Evolution toward Crown Clades. Molecular biology and evolution, 23(9), 1792–1800. 10.1093/molbev/msl044

Janssen, R., Bakker, J., & Gommers, F. (1991). Mendelian proof for a gene-for-gene relationship between virulence of Globodera tuberosum ssp. andigena CPC 1673. Rev. Nematol, 14, 207–211.

Jenkins, W. (1964). A rapid centrifugal-flotation technique for separating nematodes from soil. Plant disease reporter, 48(9).

Jones, J. T., Haegeman, A., Danchin, E. G. J., Gaur, H. S., Helder, J., Jones, M. G. K., Kikuchi, T., Manzanilla-López, R., Palomares-Rius, J. E., Wesemael, W. M. L., & Perry, R. N. (2013). Top 10 plant-parasitic nematodes in molecular plant pathology. Molecular Plant Pathology, 14(9), 946–961. 10.1111/mpp.12057

Kassambara, A. (2018). ggpubr:’ggplot2’based publication ready plots. R package version, 2.

Killick, R., & Eckley, I. A. (2014). changepoint: An R Package for Changepoint Analysis. Journal of statistical software, 58(3), 1–19. 10.18637/jss.v058.i03

Kim, D., Paggi, J. M., Park, C., Bennett, C., & Salzberg, S. L. (2019). Graph-based genome alignment and genotyping with HISAT2 and HISAT-genotype. Nature Biotechnology, 37(8), 907–915. 10.1038/s41587-019-0201-4

Koren, S., Walenz, B. P., Berlin, K., Miller, J. R., Bergman, N. H., & Phillippy, A. M. (2017). Canu: scalable and accurate long-read assembly via adaptive k-mer weighting and repeat separation. Genome research, 27(5), 722–736. 10.1101/gr.215087.116

Kriventseva, E. V., Kuznetsov, D., Tegenfeldt, F., Manni, M., Dias, R., Simão, F. A., & Zdobnov, E. M. (2018). OrthoDB v10: sampling the diversity of animal, plant, fungal, protist, bacterial and viral genomes for evolutionary and functional annotations of orthologs. Nucleic Acids Research, 47(D1), D807–D811. 10.1093/nar/gky1053

Krogh, A., Larsson, B., Von Heijne, G., & Sonnhammer, E. L. (2001). Predicting transmembrane protein topology with a hidden Markov model: application to complete genomes. Journal of molecular biology, 305(3), 567–580. 10.1006/jmbi.2000.4315

Lam, K.-K., LaButti, K., Khalak, A., & Tse, D. (2015). FinisherSC: a repeat-aware tool for upgrading de novo assembly using long reads. Bioinformatics, 31(19), 3207–3209. 10.1093/bioinformatics/btv280

Lechevalier, O., Gazengel, K., Esquibet, M., Fournet, S., Grenier, E., Daval, S., & Montarry, J. (2025). Identification through a transcriptomic approach of candidate genes involved in the adaptation of the cyst nematode Globodera pallida to the potato resistance factor GpaVvrn. BMC Genomics, 26(1), 191. 10.1186/s12864-025-11332-3

Leuenberger, J., Esnault, F., Lebas, P. L., Fournet, S., Cann, M. P., Marhadour, S., Prodhomme, C., Pilet-Nayel, M. L., & Kerlan, M. C. (2025). Identification by GWAS of marker haplotypes relevant to breed potato for Globodera pallida resistance. Theoretical and Applied Genetics, 138(3), 52. 10.1007/s00122-024-04794-8

Livak, K. J., & Schmittgen, T. D. (2001). Analysis of Relative Gene Expression Data Using Real-Time Quantitative PCR and the 2−ΔΔCT Method. Methods, 25(4), 402–408. 10.1006/meth.2001.1262

Love, M. I., Huber, W., & Anders, S. (2014). Moderated estimation of fold change and dispersion for RNA-seq data with DESeq2. Genome biology, 15, 1–21. 10.1186/s13059-014-0550-8

Milczarek, D., Flis, B., & Przetakiewicz, A. (2011). Suitability of Molecular Markers for Selection of Potatoes Resistant to Globodera spp. American Journal of Potato Research, 88(3), 245–255. 10.1007/s12230-011-9189-0

Mitchum, M. G., Hussey, R. S., Baum, T. J., Wang, X., Elling, A. A., Wubben, M., & Davis, E. L. (2013). Nematode effector proteins: an emerging paradigm of parasitism. New Phytologist, 199(4), 879–894. 10.1111/nph.12323

Mölder, F., Jablonski, K. P., Letcher, B., Hall, M. B., Tomkins-Tinch, C. H., Sochat, V., Forster, J., Lee, S., Twardziok, S. O., & Kanitz, A. (2021). Sustainable data analysis with Snakemake. F1000R*esearch*, *10*, 33. 10.12688/f1000research.29032.2

Moya, N. D., Stevens, L., Miller, I. R., Sokol, C. E., Galindo, J. L., Bardas, A. D., Koh, E. S. H., Rozenich, J., Yeo, C., Xu, M., & Andersen, E. C. (2023). Novel and improved Caenorhabditis briggsae gene models generated by community curation. BMC Genomics, 24(1), 486. 10.1186/s12864-023-09582-0

Mwangi, J. M., Niere, B., Daub, M., Finckh, M. R., & Kiewnick, S. (2019). Reproduction of Globodera pallida on tissue culture-derived potato plants and their potential use in resistance screening process. Nematology, 21(6), 613–623. 10.1163/15685411-00003239

Niere, B., Krüssel, S., & Osmers, K. (2014). Auftreten einer außergewöhnlich virulenten Population der Kartoffelzystennematoden. Journal für Kulturpflanzen, 66(12), 426–427.

Oksanen, J., Blanchet, F. G., Kindt, R., Legendre, P., Minchin, P. R., O’hara, R., Simpson, G. L., Solymos, P., Stevens, M. H. H., & Wagner, H. (2013). Package ‘vegan’. Community ecology package, version, 2(9), 1–295.

Paysan-Lafosse, T., Blum, M., Chuguransky, S., Grego, T., Pinto, B. L., Salazar, G. A., Bileschi, M. L., Bork, P., Bridge, A., Colwell, L., Gough, J., Haft, D. H., Letunić, I., Marchler-Bauer, A., Mi, H., Natale, D. A., Orengo, C. A., Pandurangan, A. P., Rivoire, C., . . . Bateman, A. (2023). InterPro in 2022. Nucleic Acids Research, 51(D1), D418–D427. 10.1093/nar/gkac993

Phillips, M. S., & Blok, V. C. (2008). Selection for reproductive ability in Globodera pallida populations in relation to quantitative resistance from Solanum vernei and S. tuberosum ssp. andigena CPC2802. Plant Pathology, 57(3), 573–580. 10.1111/j.1365-3059.2007.01771.x

Plantard, O., Picard, D., Valette, S., Scurrah, M., Grenier, E., & Mugniery, D. (2008). Origin and genetic diversity of Western European populations of the potato cyst nematode (Globodera pallida) inferred from mitochondrial sequences and microsatellite loci. Molecular Ecology, 17(9), 2208–2218. 10.1111/j.1365-294x.2008.03718.x

Price, J. A., Coyne, D., Blok, V. C., & Jones, J. T. (2021). Potato cyst nematodes Globodera rostochiensis and G. pallida. Molecular Plant Pathology, 22(5), 495–507. 10.1111/mpp.13047

R Core Team, R. (2013). R: A language and environment for statistical computing.

Rice, S., Stone, A., & leadbeater, B. (1987). Changes in cell structure in roots of resistant potatoes parasitized by potato cyst nematodes. 2. Potatoes with resistance derived from Solanum vernei. Physiological and Molecular Plant Pathology, 31(1), 1–14. 10.1016/0885-5765(87)90002-6

Roach, M. J., Schmidt, S. A., & Borneman, A. R. (2018). Purge Haplotigs: allelic contig reassignment for third-gen diploid genome assemblies. BMC bioinformatics, 19(1), 1–10. 10.1186/s12859-018-2485-7

Rouppe Van der Voort, J., Van der Vossen, E., Bakker, E., Overmars, H., Van Zandvoort, P., Hutten, R., Klein Lankhorst, R., & Bakker, J. (2000). Two additive QTLs conferring broad-spectrum resistance in potato to Globodera pallida are localized on resistance gene clusters. Theoretical and Applied Genetics, 101(7), 1122–1130. 10.1007/s001220051588

Ruan, J., & Li, H. (2020). Fast and accurate long-read assembly with wtdbg2. Nature Methods, 17(2), 155–158. 10.1038/s41592-019-0669-3

Sabeh, M., Duceppe, M.-O., St-Arnaud, M., & Mimee, B. (2018). Transcriptome-wide selection of a reliable set of reference genes for gene expression studies in potato cyst nematodes (Globodera spp.). PLOS ONE, 13(3), e0193840. 10.1371/journal.pone.0193840

Sattarzadeh, A., Achenbach, U., Lübeck, J., Strahwald, J., Tacke, E., Hofferbert, H.-R., Rothsteyn, T., & Gebhardt, C. (2006). Single nucleotide polymorphism (SNP) genotyping as basis for developing a PCR-based marker highly diagnostic for potato varieties with high resistance to Globodera pallida pathotype Pa2/3. Molecular Breeding, 18(4), 301–312. 10.1007/s11032-006-9026-1

Schouten, H. J. (1993). Models of incomplete selection for virulence of potato cyst nematodes caused by sex determination that depends on host resistance. Netherlands Journal of Plant Pathology, 99, 191–200. 10.1007/BF03041409

Schouten, H. J., & Beniers, J. E. (1997). Durability of resistance to Globodera pallida I. Changes in pathogenicity, virulence, and aggressiveness during reproduction on partially resistant potato cultivars. Phytopathology, 87(8), 862–867. 10.1094/phyto.1997.87.8.862

Sharma, S. K., Bolser, D., de Boer, J., Sønderkær, M., Amoros, W., Carboni, M. F., D’Ambrosio, J. M., de la Cruz, G., Di Genova, A., Douches, D. S., Eguiluz, M., Guo, X., Guzman, F., Hackett, C. A., Hamilton, J. P., Li, G., Li, Y., Lozano, R., Maass, A., . . . Bryan, G. J. (2013). Construction of Reference Chromosome-Scale Pseudomolecules for Potato: Integrating the Potato Genome with Genetic and Physical Maps. G3 Genes|Genomes|Genetics, 3(11), 2031-2047. 10.1534/g3.113.007153

Team, R. (2015). RStudio: integrated development for R.

Teufel, F., Almagro Armenteros, J. J., Johansen, A. R., Gíslason, M. H., Pihl, S. I., Tsirigos, K. D., Winther, O., Brunak, S., von Heijne, G., & Nielsen, H. (2022). SignalP 6.0 predicts all five types of signal peptides using protein language models. Nature Biotechnology. 10.1038/s41587-021-01156-3

Van Berloo, R., Hutten, R., Van Eck, H., & Visser, R. (2007). An online potato pedigree database resource. Potato Research, 50(1), 45–57. 10.1007/s11540-007-9028-3

van der Voort, J. R., Lindeman, W., Folkertsma, R., Hutten, R., Overmars, H., Van der Vossen, E., Jacobsen, E., & Bakker, J. (1998). A QTL for broad-spectrum resistance to cyst nematode species (Globodera spp.) maps to a resistance gene cluster in potato. Theoretical and Applied Genetics, 96(5), 654–661. 10.1007/s001220050785

van Eck, H. J., Vos, P. G., Valkonen, J. P. T., Uitdewilligen, J. G. A. M. L., Lensing, H., de Vetten, N., & Visser, R. G. F. (2017). Graphical genotyping as a method to map Ny(o,n)stoand Gpa5 using a reference panel of tetraploid potato cultivars. Theoretical and Applied Genetics, 130(3), 515–528. 10.1007/s00122-016-2831-y

van Steenbrugge, J. J. M., van den Elsen, S., Holterman, M., Lozano-Torres, Jose L., Putker, V., Thorpe, P., Goverse, A., Sterken, Mark G., Smant, G., & Helder, J. (2023). Comparative genomics among cyst nematodes reveals distinct evolutionary histories among effector families and an irregular distribution of effector-associated promoter motifs. Molecular Ecology, n/a(n/a). 10.1111/mec.16505

Varypatakis, K., Jones, J. T., & Blok, V. C. (2019). Susceptibility of potato varieties to populations of Globodera pallida selected for increased virulence. Nematology, 21(9), 995–998. 10.1163/15685411-00003283

Varypatakis, K., Véronneau, P.-Y., Thorpe, P., Cock, P. J., Lim, J. T.-Y., Armstrong, M. R., Janakowski, S., Sobczak, M., Hein, I., & Mimee, B. (2020). The Genomic Impact of Selection for Virulence against Resistance in the Potato Cyst Nematode, Globodera pallida. Genes, 11(12), 1429. 10.3390/genes11121429

Vasimuddin, M., Misra, S., Li, H., & Aluru, S. (2019). Efficient architecture-aware acceleration of BWA-MEM for multicore systems. 2019 IEEE international parallel and distributed processing symposium (IPDPS), https://doi.ieeecomputersociety.org/10.1109/IPDPS.2019.00041

Vos, P. G., Uitdewilligen, J. G., Voorrips, R. E., Visser, R. G., & van Eck, H. J. (2015). Development and analysis of a 20K SNP array for potato (Solanum tuberosum): an insight into the breeding history. Theoretical and Applied Genetics, 128, 2387–2401. 10.1007/s00122-015-2593-y

Walker, B. J., Abeel, T., Shea, T., Priest, M., Abouelliel, A., Sakthikumar, S., Cuomo, C. A., Zeng, Q., Wortman, J., & Young, S. K. (2014). Pilon: an integrated tool for comprehensive microbial variant detection and genome assembly improvement. PLOS ONE, 9(11), e112963. 10.1371/journal.pone.0112963

Wang, Y., Brown, L. H., Adams, T. M., Cheung, Y. W., Li, J., Young, V., Todd, D. T., Armstrong, M. R., Neugebauer, K., Kaur, A., Harrower, B., Oome, S., Wang, X., Bayer, M., & Hein, I. (2023). SMRT–AgRenSeq-d in potato (Solanum tuberosum) as a method to identify candidates for the nematode resistance Gpa5. Horticulture Research, 10(11). 10.1093/hr/uhad211

Xu, X., Pan, S., Cheng, S., Zhang, B., Mu, D., Ni, P., Zhang, G., Yang, S., Li, R., Wang, J., Orjeda, G., Guzman, F., Torres, M., Lozano, R., Ponce, O., Martinez, D., De la Cruz, G., Chakrabarti, S. K., Patil, V. U., . . . Research, C. (2011). Genome sequence and analysis of the tuber crop potato. Nature, 475(7355), 189–195. 10.1038/nature10158

Zheng, Q., Bertran, A., Brand, A., Van Schaik, C. C., van de Ruitenbeek, S. J., Smant, G., Goverse, A., & Sterken, M. G. (2022). Comparative Transcriptome Analysis Reveals the Specific Activation of Defense Pathways Against Globodera pallida in Gpa2 Resistant Potato Roots. Frontiers in Plant Science, 13. 10.3389/fpls.2022.909593

