## Supplementary figures and images for "The potato cyst nematode *Globodera pallida* overcomes major potato resistance through selection on standing variation at a single locus"

### Supplementary figure 1

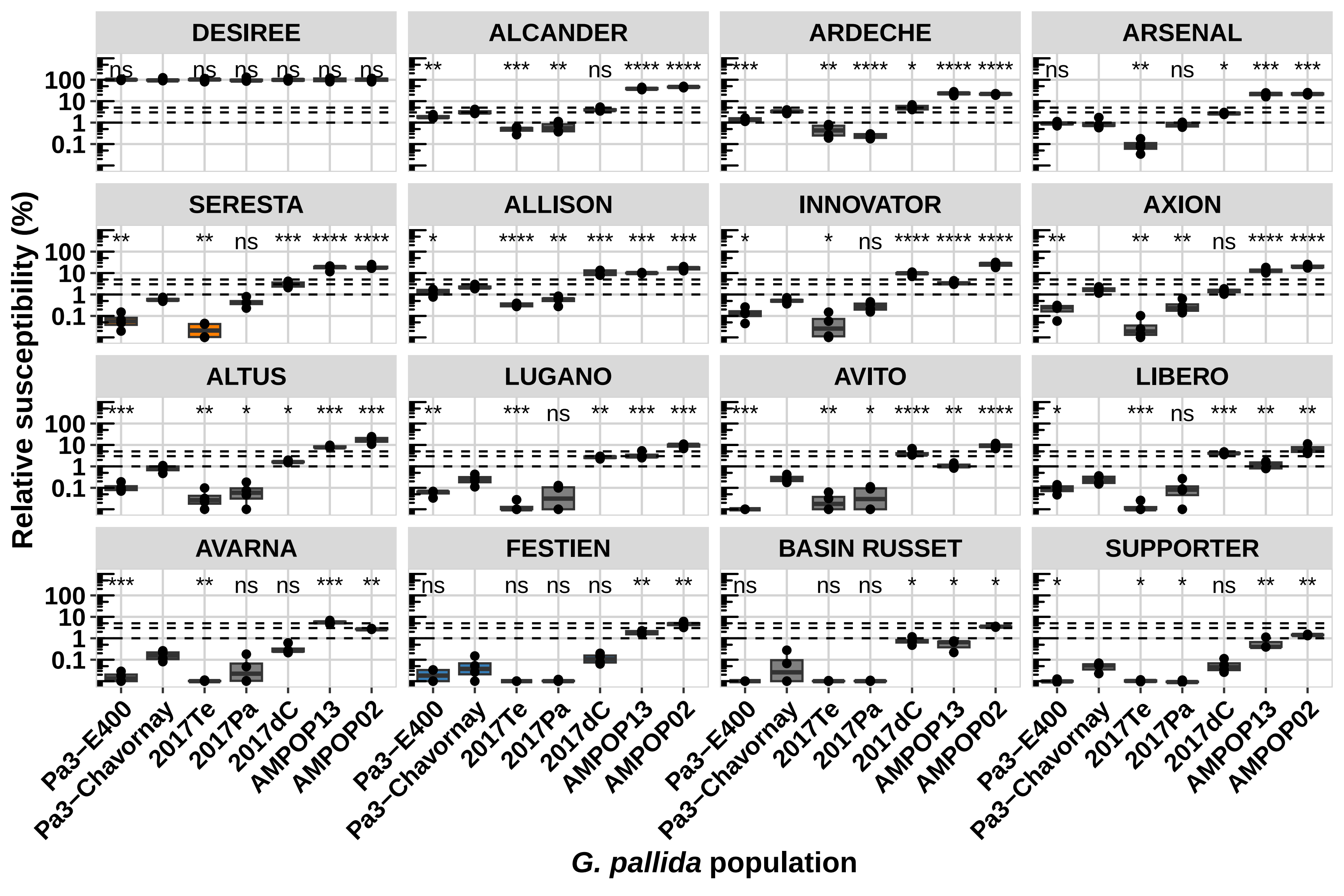

### Supplementary figure 2

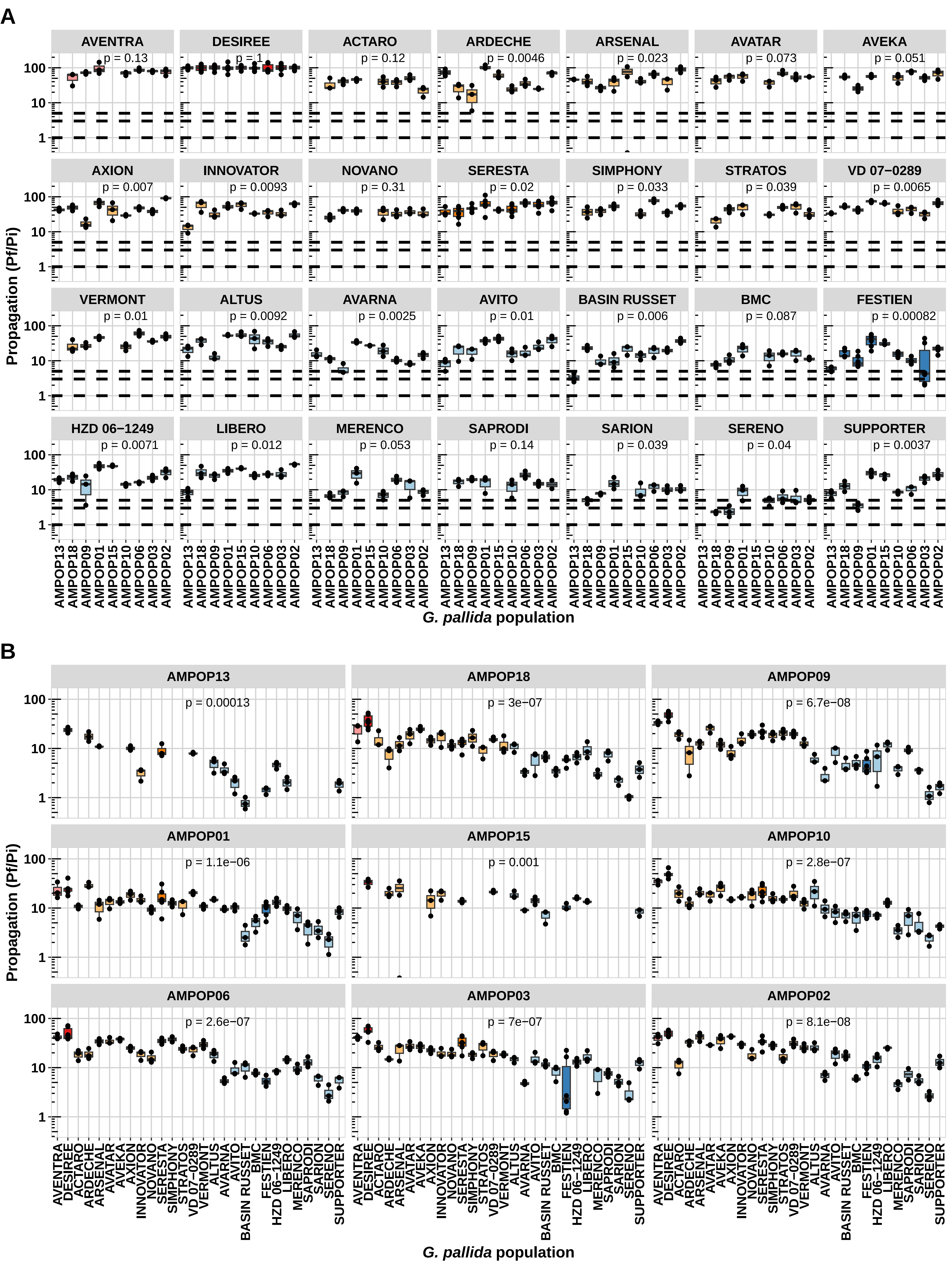

### Supplementary figure 3

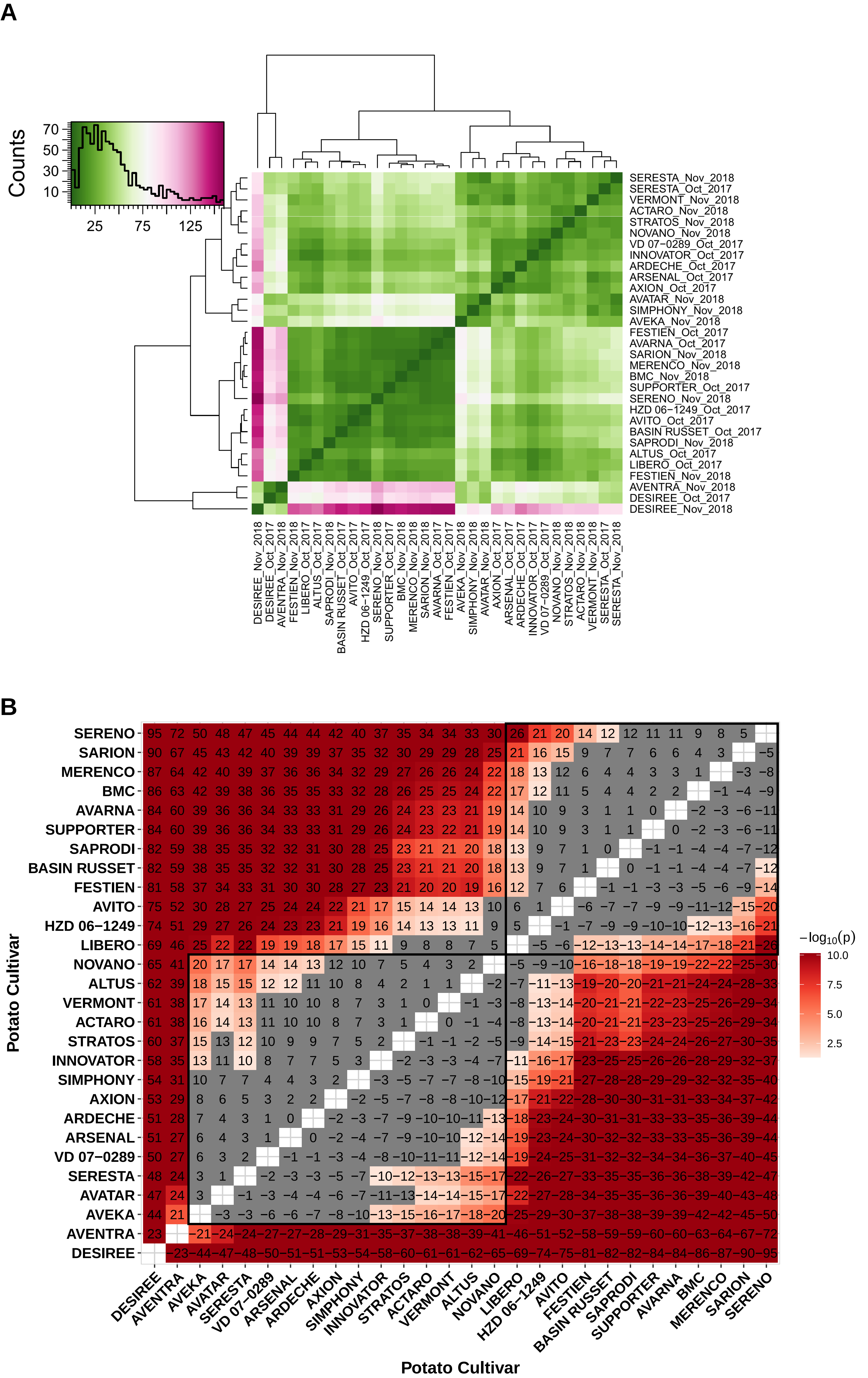

### Supplementary figure 4

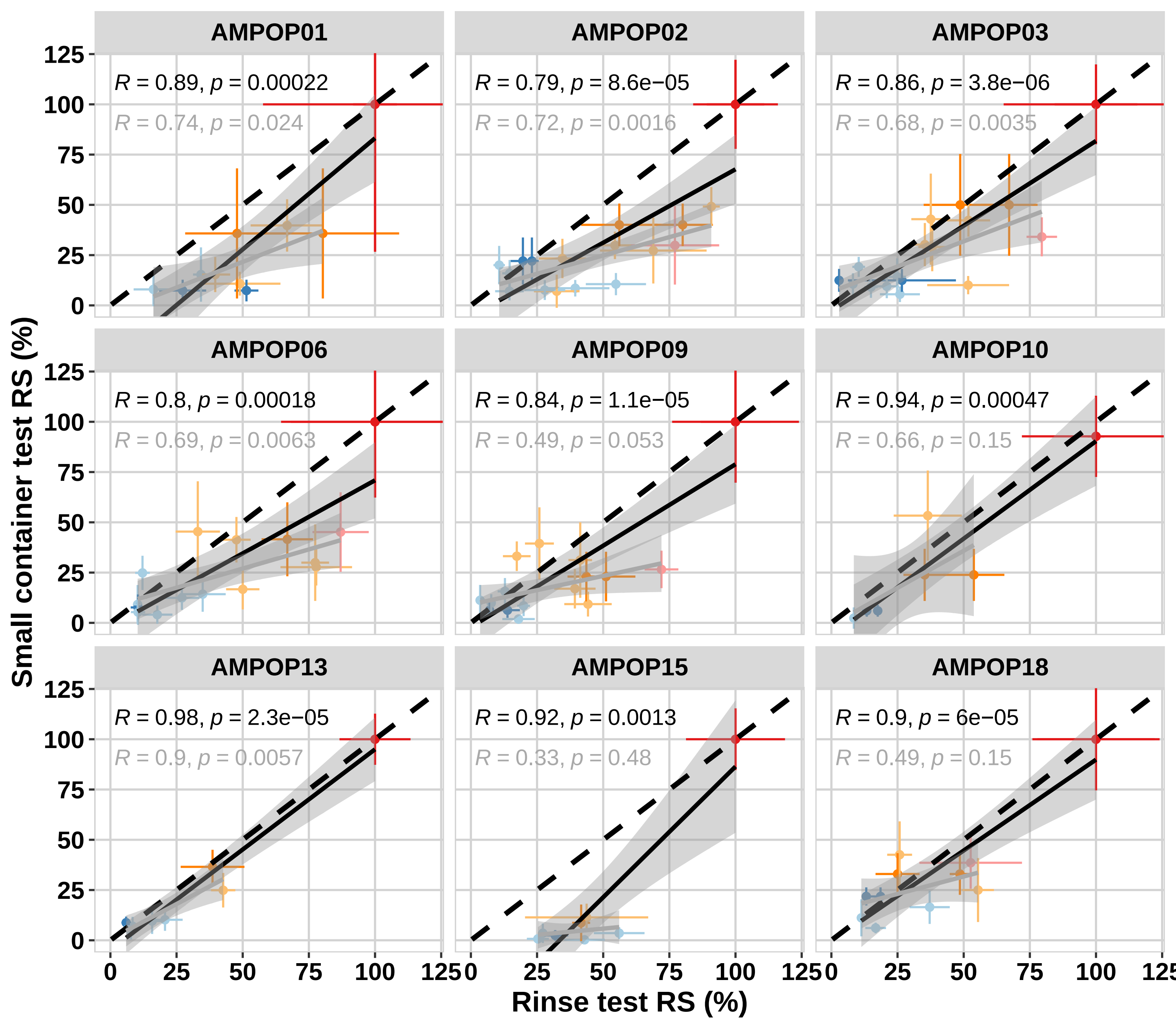

### Supplementary figure 5

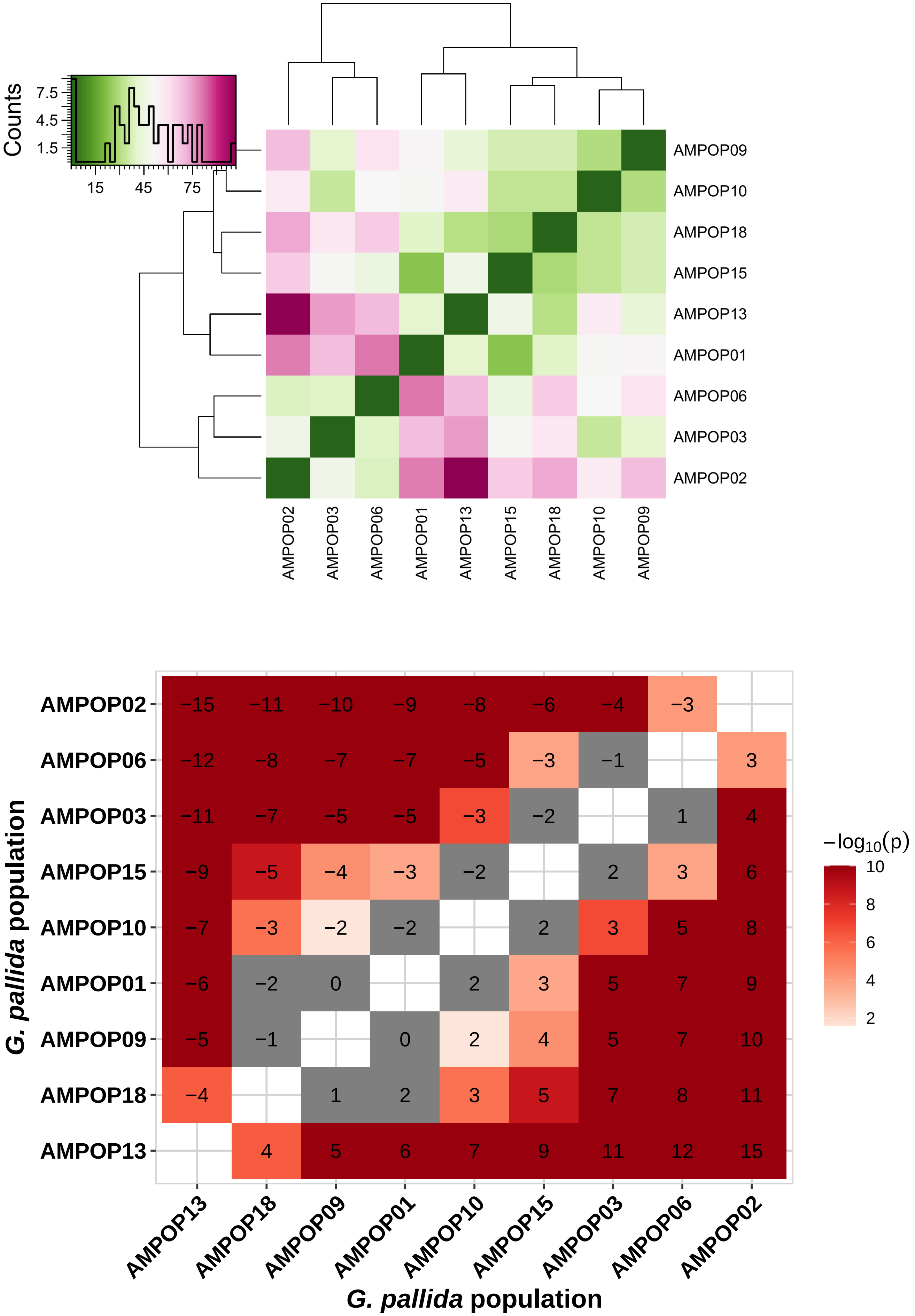

### Supplementary figure 6

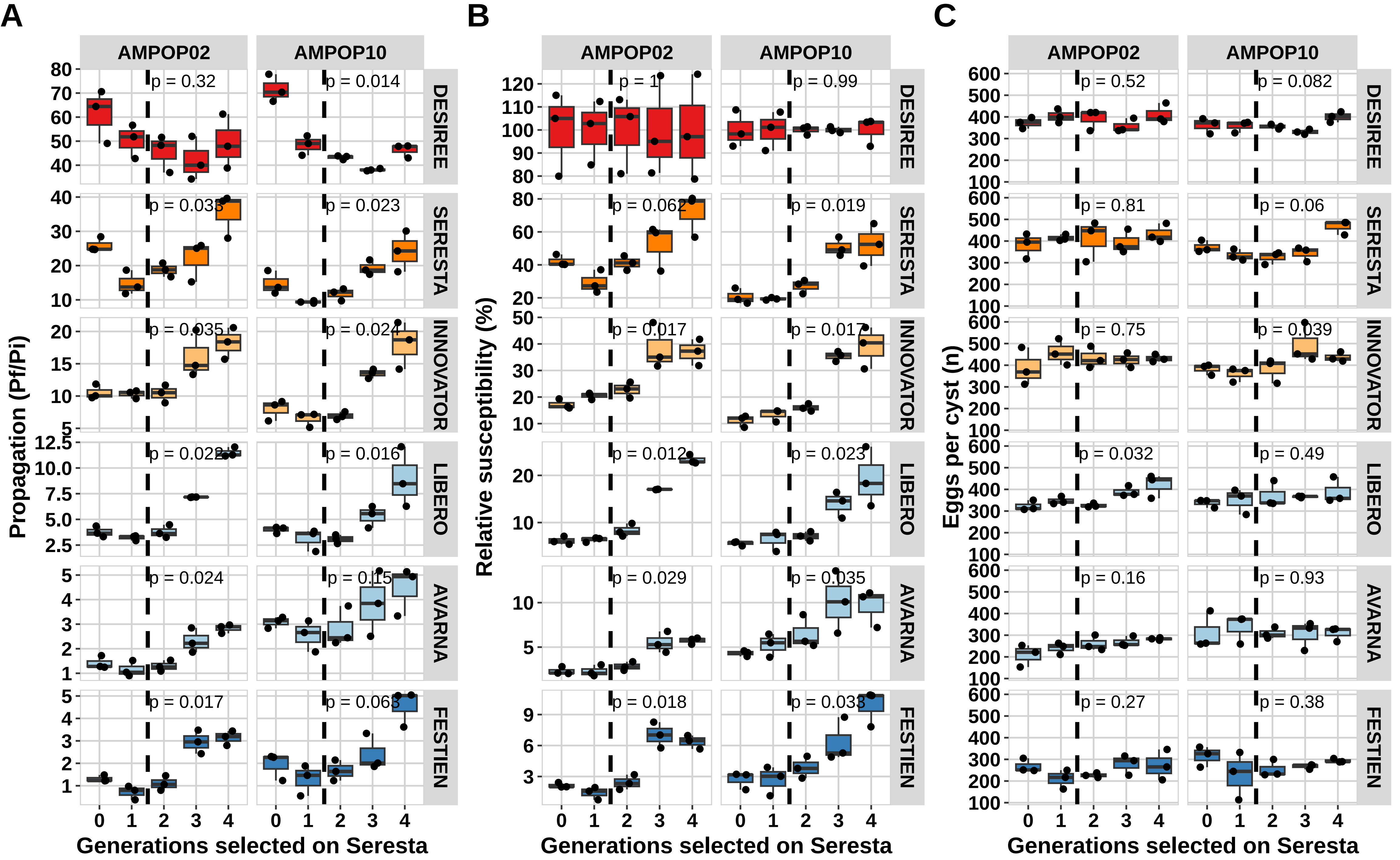

### Supplementary figure 7

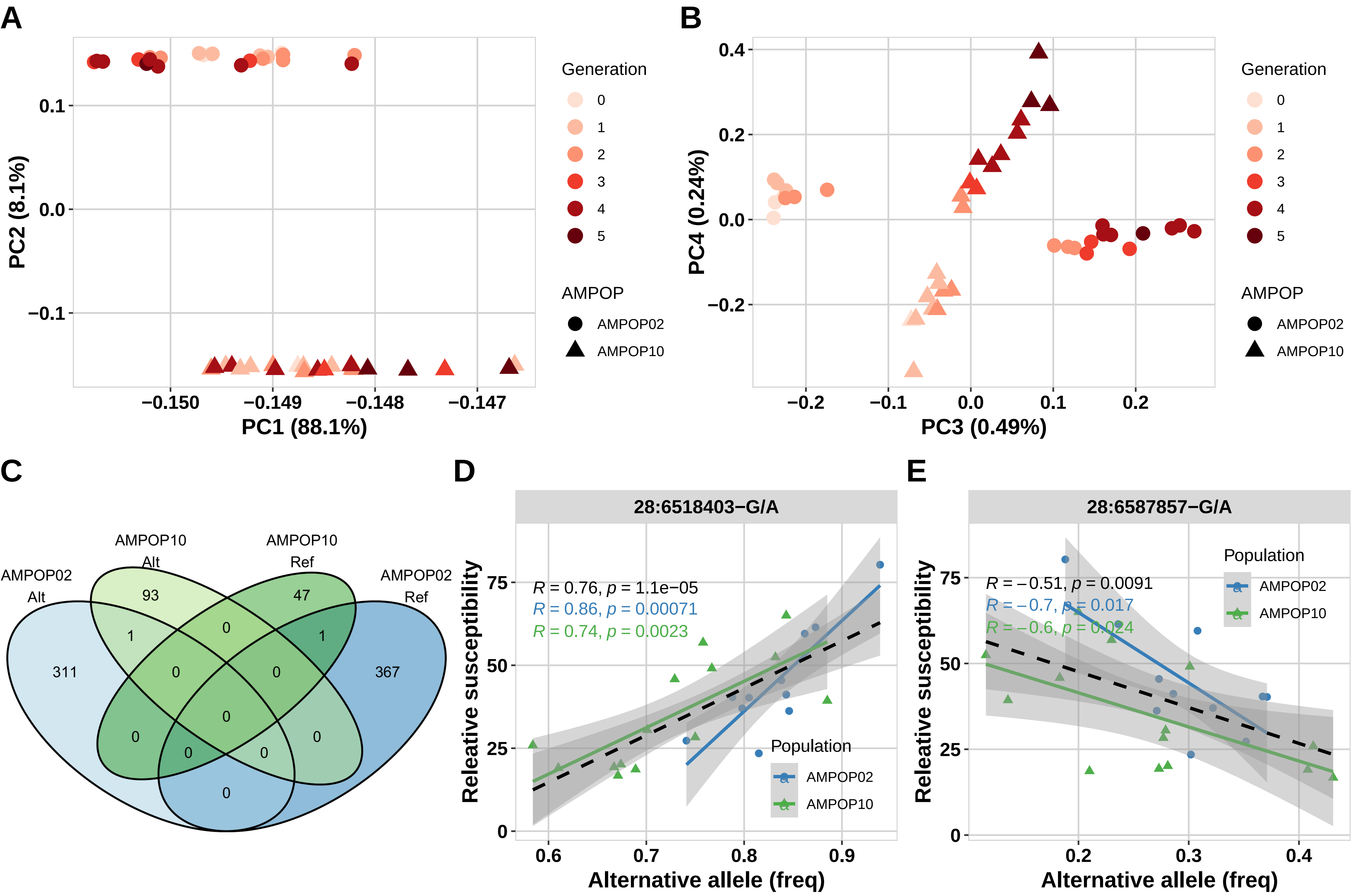

### Supplementary figure 8

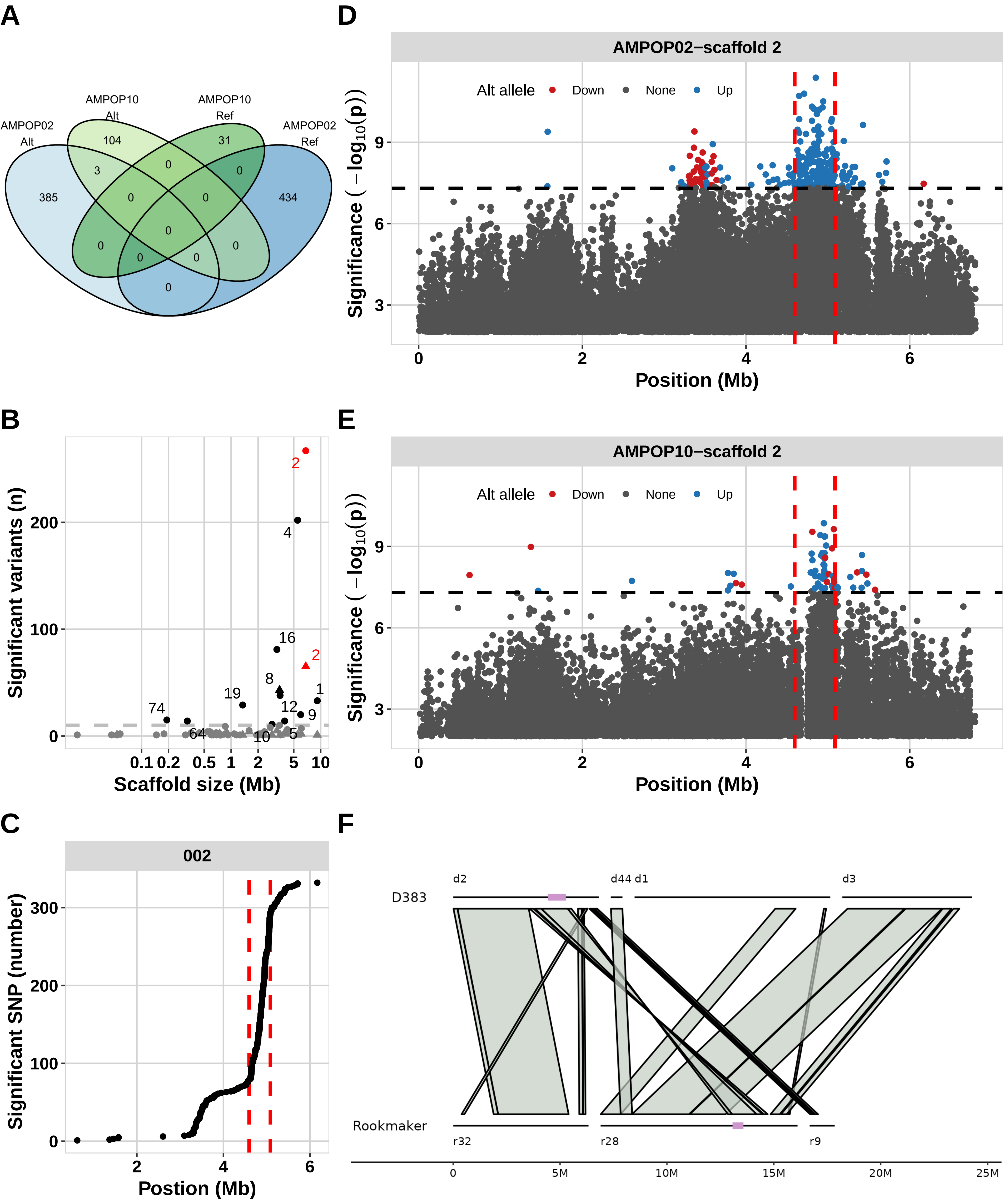

### Supplementary figure 9

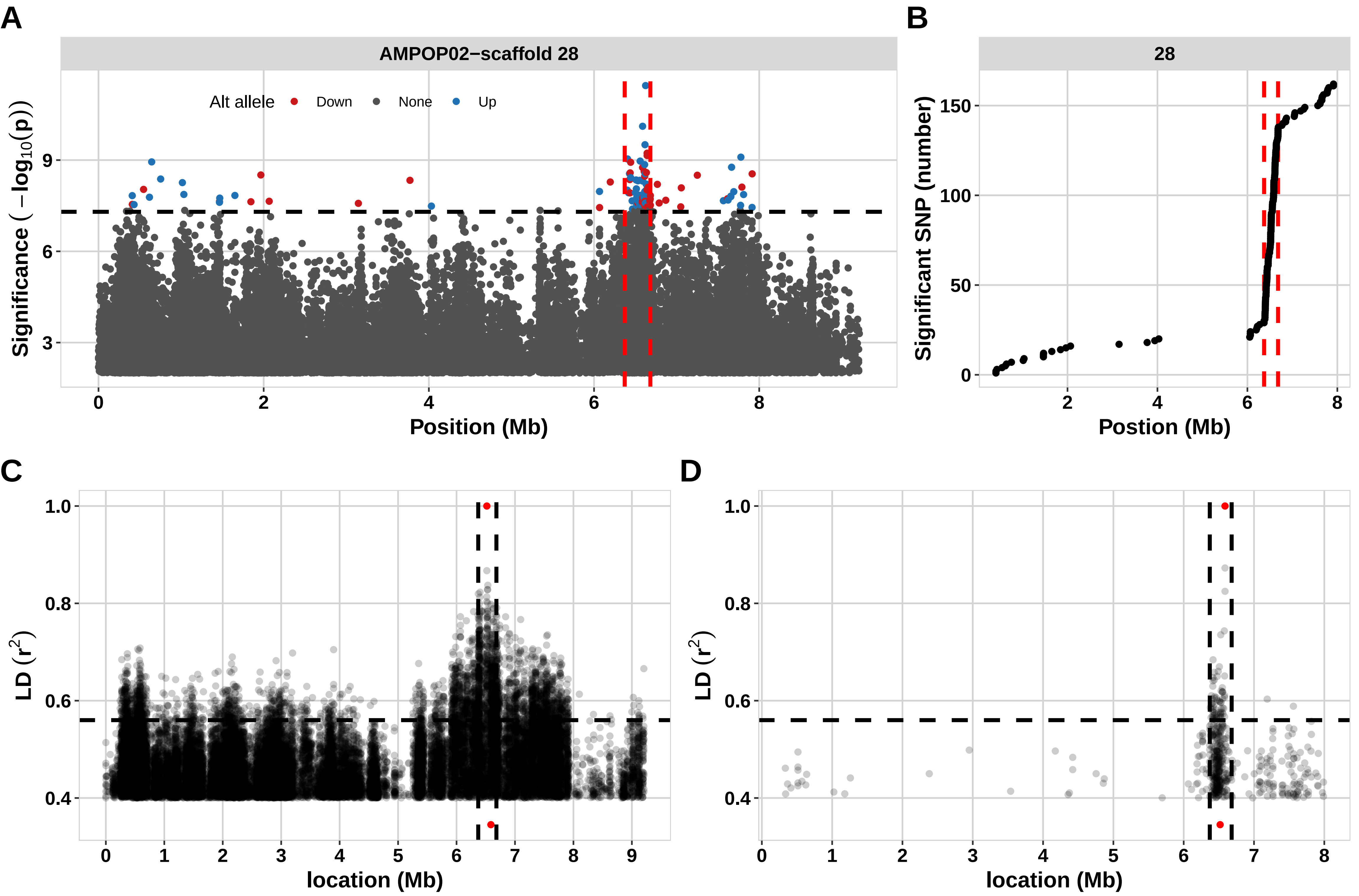
